# A primed immune transcriptional program is activated in oligodendroglia in multiple sclerosis

**DOI:** 10.1101/2020.07.21.213876

**Authors:** Mandy Meijer, Eneritz Agirre, Mukund Kabbe, Cassandra A. van Tuijn, Abeer Heskol, Ana Mendanha Falcão, M. Ryan Corces, Thomas J. Montine, Xingqi Chen, Howard Y. Chang, Gonçalo Castelo-Branco

## Abstract

Multiple sclerosis (MS) is a disease characterized by a targeted immune attack on myelin in the central nervous system (CNS). We have previously shown that oligodendrocytes (OLs), myelin producing cells in the CNS, and their precursors (OPCs), acquire disease-specific transcriptional states in MS ^1,2^. To understand how these alternative transcriptional programs are activated in disease, we performed single-cell assay for transposase accessible chromatin using sequencing (scATAC-seq) on the OL lineage in the experimental autoimmune encephalomyelitis (EAE) mouse model of MS. We identified regulatory regions with increased accessibility in oligodendroglia (OLG) in EAE, some of which in the proximity of immune genes. A similar re-modeling of chromatin accessibility was observed upon treatment of postnatal OPCs with interferon-gamma (IFN*γ*), but not with dexamethasone. These changes in accessibility were not exclusive to distal enhancers, but also occurred at promoter regions, suggesting a role for promoters in mediating cell-state transitions. Notably, we found that a subset of immune genes already exhibited chromatin accessibility in OPCs ex vivo and in vivo, suggesting a primed chromatin state in OLG compatible with rapid transitions to an immune-competent state. Several primed genes presented bivalency of H3K4me3 and H3K27me3 at promoters in OPCs, with loss of H3K27me3 upon IFN*γ* treatment. Inhibition of JMJD3/*Kdm6b*, mediating removal of H3K27me3, led to the inability to activate these genes upon IFN*γ* treatment. Importantly, OLGs from the adult human brain showed chromatin accessibility at immune gene loci, particularly at MHC-I pathway genes. A subset of single-nucleotide polymorphisms (SNPs) associated with MS susceptibility overlapped with these primed regulatory regions in OLG from both mouse and human CNS. Our data suggest that susceptibility for MS may involve activation of immune gene programs in OLG. These programs are under tight control at the chromatin level in OLG and may therefore constitute novel targets for immunological-based therapies for MS.

## Introduction

OLs facilitate neuronal communication in the CNS by coordinating axonal myelination. These myelin sheaths are targeted by the immune system in MS. While OPCs in the adult CNS have been thought to be recruited to MS lesions and contribute to remyelination during disease remission, this capacity might be hindered upon disease progression ^2,3^. We and others have shown that OPCs transition to an immune-like state in the context of MS and demyelination ^1,2,4,5^, characterized by the upregulation of immune response genes, including major histocompatibility complex (MHC)-I and MHC-II. In this state, immune OPCs can present antigens to both CD4^1^ and CD8 cytotoxic T-cells ^4,5^. This transition diminishes the ability of immune OPCs to differentiate into mature OLs ^4^, which might compromise remyelination.

Single-nucleus RNA sequencing of white matter of MS patients suggests that a population of OLs is also induced to express immune genes ^2,6^. Nevertheless, phagocytosing microglia/macrophages have also been suggested to import myelin transcripts into their nucleus ^6^, which could blur the proper distinction between immune cells and OLG, and thus constitute a challenge in the determination of cell identities. To resolve if indeed immune transcriptional programs can be operational in OLG and to investigate the mechanisms by which they are established, we profiled the chromatin accessibility of OLG at the single-cell level in the EAE mouse model of MS.

### Differential chromatin accessibility allows identification of disease-specific OLG in the EAE mouse model of MS

We performed scATAC-seq on the spinal cords of Sox10:Cre-RCE:LoxP(EGFP) mice ^1,7,8^ induced with MOG35-55 peptide in CFA (EAE) or with CFA alone (CFA-Ctr) (Fig. 1a). Spinal cords were isolated at the peak of the disease (when EAE animals reached score 3), freshly dissociated GFP+ (labeling OLG) and GFP-cells were sorted and pooled 4:1. The pool of cells was then used as input for the 10x Genomics scATAC-seq protocol (Fig. 1a). Clustering considering genome wide differences in chromatin accessibility allowed the identification of 20 different clusters (Fig. 1b). Accessibility at the *Sox10* locus was detected in all clusters except clusters 4, 10, 12, 15, 16, 17 and 18 (Supplementary Fig. 1a), from which 4, 10, 15 and 17 presented accessibility at the *Aif1* locus (marker for microglia and related cells, which might also include macrophages (MiGl)). Accessibility at the *Ptprz1* (marker of OPCs) and *Mog* (marker of mature OL, MOL) loci was detected in smaller distinct subsets of clusters presenting also accessibility at *Sox10* (Supplementary Fig. 1a). Notably, both OPC and MOL populations derived from EAE mice clustered separately from CFA-Ctr animals, while MiGl came exclusively from EAE (Fig. 1c, d).

**Fig. 1.**
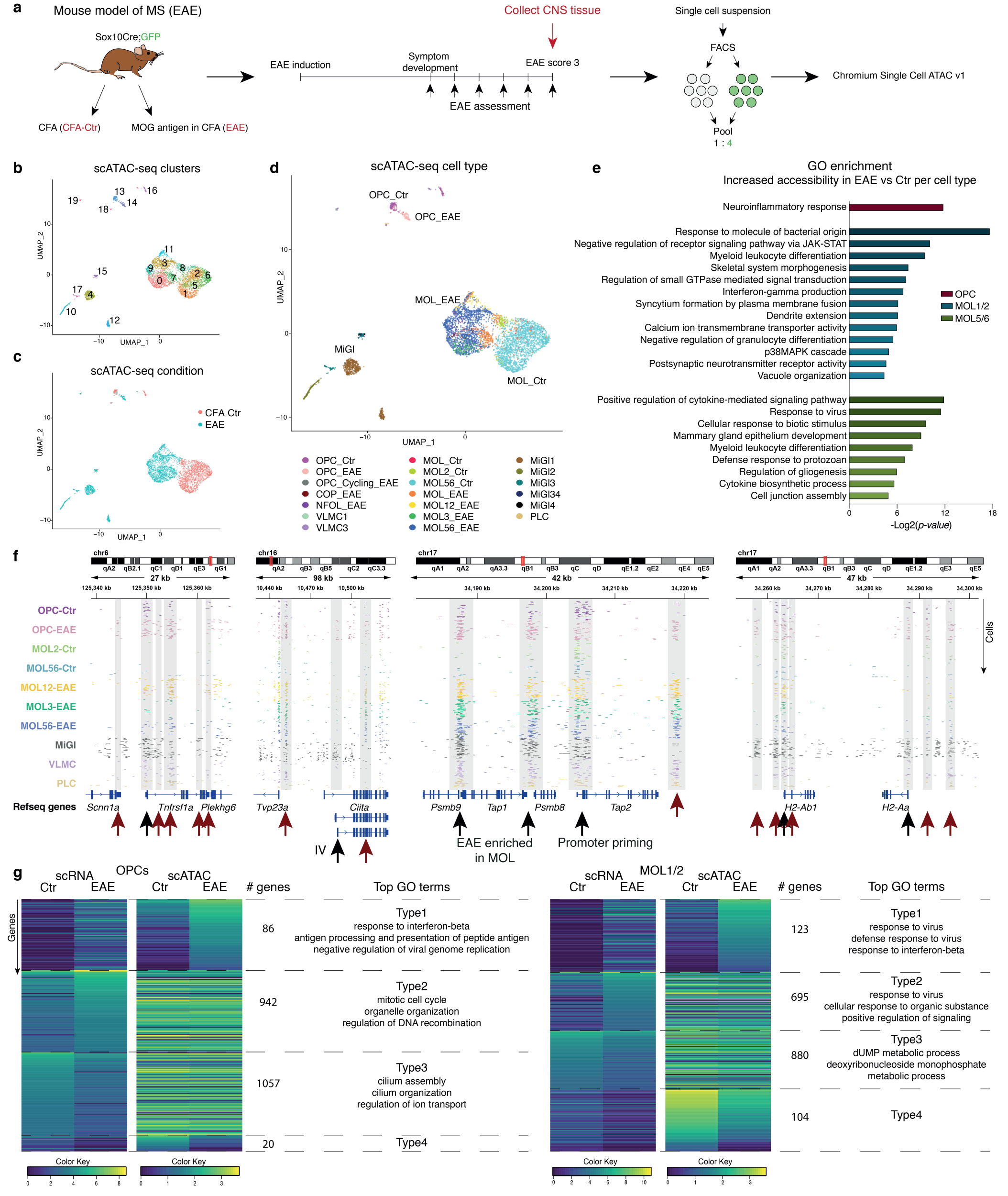
Single-cell ATAC-seq reveals primed and increased chromatin accessibility at immune gene loci in oligodendroglia from the EAE mouse model of MS. **a**, Schematic illustration of the experimental setup for scATAC-seq of oligodendroglia from the EAE mouse model of MS. **b**, UMAP clustering based on chromatin accessibility reveals 20 different clusters. **c**, Experimental condition origin of individual cells, projected on top of UMAP clustering. **d**, Label transfer from previous matched scRNA-seq data 1 projected on top of UMAP clustering. **e**, Gene Ontology enrichment analysis (biological processes) of the nearest genes to enriched accessibility peaks in EAE are shown for OPC, MOL1/2 and MOL5/6 populations. **f**, Integrative Genomics Viewer (IGV) tracks of chromatin accessibility in 50 randomly selected individual cells for each selected cluster, with MiGl clusters grouped together. Highlighted with grey boxes and arrows are regions with differential accessibility in specific clusters or promoter priming, black arrows indicate promoter regions and red arrows indicate putative enhancer. Genomic coordinates are shown. **g**, Genes in OPCs (left) or MOL1/2 (right) are clustered based on gene expression differences between EAE vs. Ctr and correlation with chromatin accessibility activity score (accessibility over 500 bp promoter region). Top GO terms are shown for Type1 (genes with increased expression in EAE and chromatin accessibility), Type2 (genes with increased expression in EAE, but no change in chromatin accessibility) and Type3 (genes with reduced expression in EAE, but no change in chromatin accessibility). Type4 (genes with reduced expression and chromatin accessibility in EAE) had no GO terms because of too few genes in this group.

Chromatin accessibility was observed mainly in intergenic (26.43%) and intronic regions (48.83%) (Supplementary Fig. 1b). Cell-type specific expression has been mainly associated with distal enhancers, while promoters have been suggested to provide limited information due to similar chromatin signatures across cell types ^9,10^. Nevertheless, we observed 13.84% of accessibility peaks at promoter regions (Supplementary Fig. 1b). We thus integrated EAE scRNA-seq data1 (UMAP visualization in Supplementary Fig. 1c) to the scATAC-seq gene activities over promoter regions, to classify cell types by identifying shared correlation patterns (Seurat, Signac) ^11^. We could successfully classify MiGl, vascular leptomeningeal cells (VLMCs), pericyte-like cells (PLCs), OPCs and different MOL populations, segregated as expected between CFA-Ctr and EAE (Fig. 1d, Supplementary Fig. 1d for prediction scores). However, a small subset of cells from EAE mice were classified as Ctr MOL populations and vice-versa, revealing a certain degree of ambiguity in the promoter-based scATAC-seq classification score (Supplementary Fig. 1d). These cells were manually corrected as MOL-EAE and MOL-Ctr (Fig. 1d) and excluded from further analysis. Similar results were obtained with a plate-based scATAC-seq protocol (Pi-ATAC) ^12^ on fixed cells derived from CFA-Ctr and EAE mouse brains and spinal cords (Supplementary Fig. 1e). Thus, these results indicate that OLG derived from EAE animals have an altered chromatin accessibility state and that chromatin accessibility at promoters provides relevant information to predict cell-types and possibly cell states.

### Increased chromatin accessibility at promoters and enhancers of immune genes in single OPCs and MOLs in EAE mice

To investigate what is underlying these differences in chromatin accessibility between CFA-Ctr and EAE mice, we inspected which classes of genes are closest to peaks differentially accessible in different OLG populations, by performing gene ontology (GO)-analysis (Supplementary Table 1). The nearest genes to higher accessible regions in EAE-OPCs when compared to Ctr-OPCs were involved in ‘neuroinflammatory response’, while in MOL1/2-EAE compared to MOL2-Ctr the top terms were ‘response to molecule of bacterial origin’, ‘negative regulation of receptor signaling pathway via JAK-STAT’ and ‘myeloid leukocyte differentiation’ (Fig.1e, Supplementary Table 1). For MOL5/6-EAE compared to MOL5/6-Ctr we found ‘positive regulation of cytokine-mediated signaling path-way’, ‘response to virus’ and ‘cellular response to biotic stimulus’ (Fig. 1e, Supplementary Table 1). These GO-terms indicate that chromatin accessibility changes in EAE-OLG populations are related to immune pathways, which is consistent with previously shown disease states of OLG in EAE characterized by the induction or elevated expression of genes involved in immune processes ^1^.

**Table 1.** Primers sequences for qRT-PCR.

| Primers | Forward 5' → 3' | Reverse 5' → 3' |
| --- | --- | --- |
| *Ubc* | agg tca aac agg aag aca gac gta | tca cac cca aga aca agc aca |
| *b-Act* | ctc tgg ctc cta gca cca tga aga | gta aaa cgc agc tca gta aca gtc cg |
| *Nlrc5* | ccg tgg tac tca cat ttg cc | cct tcg aga tct ctg gga ca |
| *Psmb8* | cct tac ctg ctt ggc acc at | atg ctg cag aca cgg aga |
| *H2-q7* | ata cct gca gct cgg gaa g | agc acc ata aga cct ggg gt |
| *Ciita* | ctg gca cag gtc tct cca gt | tac tga ggc tgc ttg aag gg |
| *H2-aa* | gct cag cct ctg tgg agg | tgt act ggc caa tgt ctc ca |
| *H2-ab1* | gca gac aca act acg agg gg | agt gtt gtg gtg gtt gag gg |

We observed that 11.48% of differential accessibility peaks between OLG from EAE and CFA-Ctr mice were at promoter regions (Supplementary Fig. 1b). One of the genes that had increased accessibility at the promoter in both MOL1/2-EAE and MOL5/6-EAE populations is *Tnfrsf1a* (Fig. 1f), a MS susceptibility gene encoding for TNF Receptor Superfamily Member 1A ^13^, which also presented increased gene expression in EAE scRNA-seq data ^1^ (Supplementary Fig. 2a). Similar correlations could be observed at the MHC-I locus containing genes important for antigen processing (*Psmb9*/*Tap1* and *Psmb8*; Fig.1f, Supplementary Fig. 2a). *Ciita*, a master regulator of the MHC-II pathway, had EAE-OLG specific accessibility at one intronic promoter (promoter IV, which is IFN*γ* -responsive and active in non-professional antigen presenting cells ^14,15^), while MiGl had accessibility at several promoter sites (Fig. 1f).

**Fig. 2.**
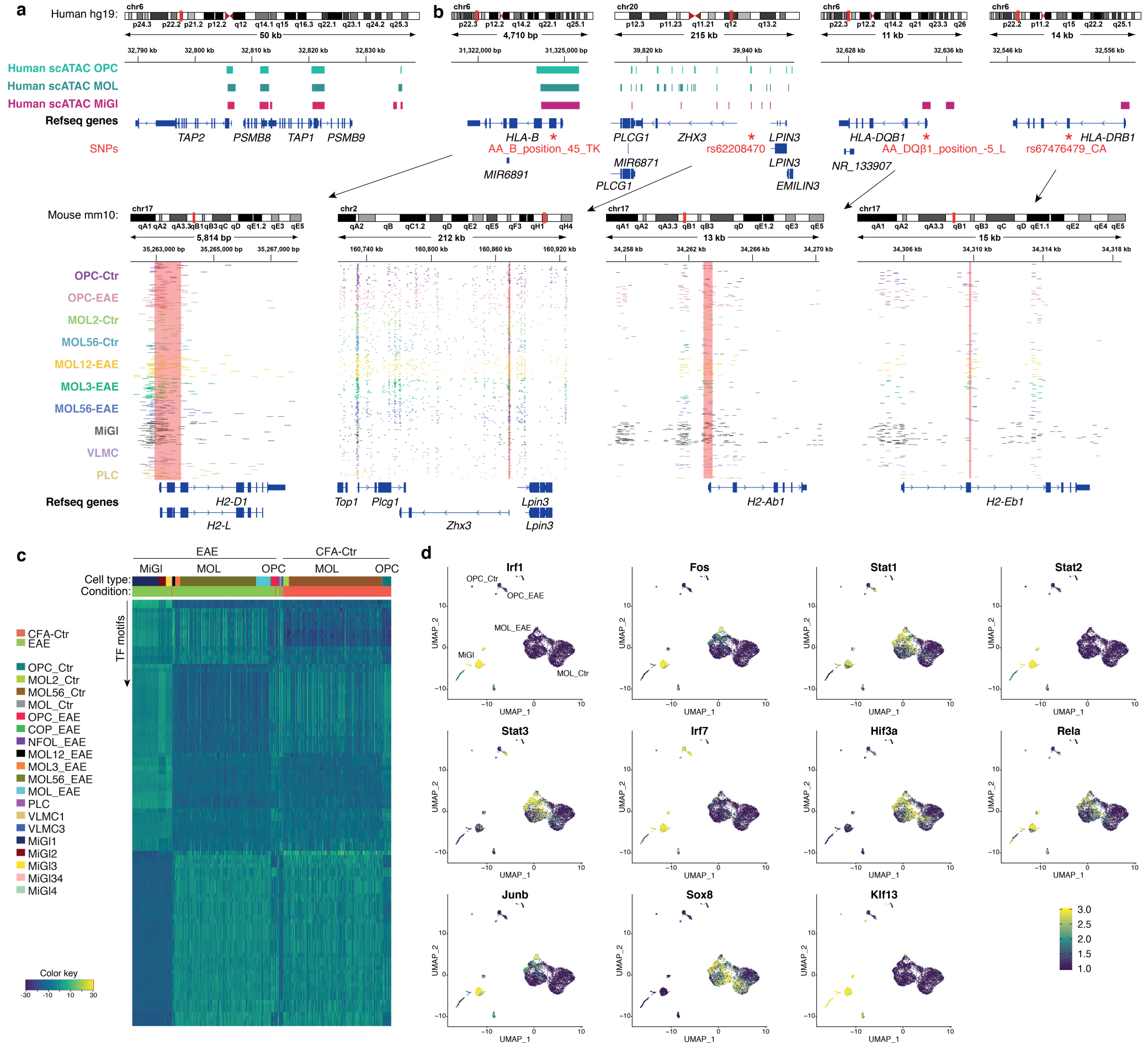
Accessible regulatory regions in oligodendroglia overlap with SNPs and outside variants conferring susceptibility in MS and have enrichment of immune transcription factor motifs in EAE. **a**, IGV tracks of chromatin accessibility regions derived from scATAC-seq populations from the adult brain from human healthy individuals are shown at MHC pathway gene loci. **b**, SNP coordinates for four MS SNPs in the hg19 human genome reference are shown with corresponding chromatin accessibility regions derived from merged scATAC-seq populations from the adult brain from human healthy individuals. Also, corresponding locations are shown in the mouse mm10 genome reference with IGV tracks of chromatin accessibility in 50 randomly selected individual cells from scATAC-seq from EAE and CFA-Ctr mice. Red boxes show scATAC-seq peaks from mouse overlapping with SNP location. **c**, ChromVAR clustering of transcription factor (TF) motif variability from scATAC-Seq. Each row presents a TF motif, while each column represents a single cell. Scale - blue (low TF motif accessibility) to yellow (high TF motif accessibility). **d**, TF motif variability projected on top of UMAP clustering of scATAC-seq.

49.02% and 33.65% of differential accessibility peaks between OLG from EAE and CFA-Ctr mice were at intergenic and intronic regions (Supplementary Fig. 1b). Distal enhancers are in contact with the transcription start site (TSS) of their target genes via long range interactions, facilitated by cell-type specific chromatin architectures ^16^. We observed chromatin accessibility at putative distal enhancers to immune genes in different EAE populations. For example, a putative enhancer at the *Tnfrsf1a* locus had accessibility only in EAE-OPCs (Fig. 1f). Other putative enhancers in the first intron of *Tnfrsf1a* showed either accessibility in MiGl and EAE-OPCs or in all EAE populations (Fig. 1f). At the *Ciita* locus we also observed specific enhancer accessibility in EAE-OLG populations within exon 10, which is not observed in MiGl (Fig. 1f). *H2-ab1*, as *Tap2*, also presented increased accessibility in EAE-OPCs and - MOLs at distal regulatory regions, but not at the TSS. An additional putative upstream enhancer of *H2-ab1* only showed chromatin accessibility in MiGl and EAE-OPCs, but not in other populations (Fig. 1f). Thus, EAE derived OLG have increased chromatin accessibility both at the promoters and enhancers of genes related to immune pathways, suggesting that modulation of chromatin at not only distal but also proximal regions can lead to transitioning to disease specific states. Different cell-types can employ different enhancers and/or promoters and can therefore provide additional cell-type specific information of disease state and point towards cell-type specific regulation of immune pathways.

### Primed chromatin accessibility of immune genes in single OPCs and MOLs

*Klk8* has previously been identified as a marker gene specific for MOL1/2-EAE ^1^ (Supplementary Fig. 2a) and indeed we found enriched promoter chromatin accessibility in this population as well (Supplementary Fig. 2b). However, *Trim34a*, also a marker for MOL1/2-EAE, had more widespread accessibility in all EAE populations (Supplementary Fig. 2a, b). *Tlr3*, another marker for MOL1/2-EAE, and *Plin4* and *Hif3a*, two marker genes for MOL5/6-EAE-a, had promoter accessibility in most of the OLG populations found with scATAC-seq, even in control populations (Supplementary Fig. 2a, b). We also observed that several immune genes, such as *Psmb9, Tap1, Tap2* and *H2-ab1*, with increased expression in EAE-OLG were already accessible in Ctr-OLG at their promoters, in particular in Ctr-OPCs (Fig. 1f, Supplementary Fig. 2a).

We thus assessed the correlation of accessibility over 500 bp promoter regions (gene activity score) with gene expression from the scRNA-seq populations. We performed GO-analysis on the genes with increased expression in EAE and that have increased accessibility as well (Type 1). The top GO-terms for OPCs (EAE vs. Ctr) were ‘response to interferon-beta’, ‘antigen processing and presentation of peptide antigen’ and ‘negative regulation of viral genome replication’ (Fig. 1g, Supplementary Table 2). For genes that increased in EAE-OPCs, but had no change in chromatin accessibility (Type2) we found GO-terms related to antigen presentation, but also ‘mitotic cell cycle’, ‘organelle organization’ and ‘regulation of DNA recombination’ (Supplementary Table 2). Likewise, genes with higher expression in MOL1/2-EAE and MOL5/6-EAE, and increased (Type1) or similar (Type2) chromatin accessibility are associated with GO-terms related to immune processes (Fig. 1g, Supplementary Fig. 2c, Supplementary Table 2). Type 2 genes present a higher level of expression than Type1 genes in both OPCs and MOLs, which might be consistent with already open chromatin at promoters. Interestingly, genes that lose expression in EAE-OPCs/MOLs, with GO-terms related to regulation of ion transport and metabolic processes, do not lose chromatin accessibility (Type3, Fig. 1g, Supplementary Fig. 2c), and genes with reduced expression that also lose chromatin accessibility are very few (Type4). Thus, Ctr-OLG are already in a primed immune chromatin state at the promoter level, with the observed increased expression of primed immune genes in EAE most likely regulated at the enhancer level or by other mechanisms regulating chromatin at promoters.

### A subset of SNPs and outside variants conferring susceptibility in MS are located at accessible regulatory regions in OLG in the mouse and human CNS

Our results indicate that while a subset of immune genes in OLG increase their expression and chromatin accessibility at promoters in the context of EAE, another subset of immune genes have already accessibility at their promoters in CFA-Ctr. We thus investigated the chromatin accessibility of these genes in scATAC-Seq data from the adult human brain from healthy individuals ^17^. Strikingly, we observed chromatin accessibility at the promoters of MHC-I pathway genes *TAP2, PSMB8, TAP1* and *PSMB9* not only in microglia, but also in OPCs and MOLs (Fig. 2a), similarly to what we observed in CFA-Ctr mice.

We had previously shown that several MS susceptibility genes are expressed in Ctr-OPCs and EAE-OLG ^1^, suggesting that not only immune cells but also OLG might contribute to the etiology of MS. We thus assessed whether we could identify putative MS susceptibility loci associated with chromatin accessibility in OLG. We first lifted over the coordinates of all Ctr and EAE scATAC-seq peaks identified in each cell population from mouse to human. We then cross-referenced this dataset with the location of MS susceptibility SNPs ^18^ and outside variants ^19^. Linkage disequilibrium score regression (LDSC) ^20^ and Multi-marker Analysis of GenoMic Annotation (MAGMA) ^21^ analysis indicated an enrichment of chromatin accessible regions and associated genes mainly in MiGl, consistent with association of MS susceptibility with gene expression in microglia ^1,18^, and with a lesser extent in OLG (Supplementary Table 3). SNPs located at the human MHC locus *HLA-B* (SNP ID AA_B_position_45_TK ^18^) and the non-MHC locus *ZHX3* (SNP ID rs62208470^18^) coincided with chromatin accessibility at the corresponding mouse loci in MiGl and in OLG in both EAE and CFA-Ctr (Fig. 2b, Supplementary Table 3). Moreover, analysis of scATAC-seq data derived from healthy individuals revealed that also these SNPs overlapped with chromatin accessibility in human OPCs, MOLs and MiGl in a non-disease context (Fig. 2b, Supplementary Table 3).

Additional MHC and non-MHC SNPs, and outside variants also exhibited chromatin accessibility in Ctr- and EAE-OLG, such as at the promoter of H2-ab1 in MiGl and in OLG in both EAE and CFA-Ctr (SNP ID AA_DQβ1_position_-5_L at *HLA-DQB1* ^18^), and at a putative enhancer region of *H2-eb1* in EAE-OLG, but interestingly not in MiGl (SNP ID rs67476479_CA at *HLA-DRB1* ^18^) (Fig. 2b, Supplementary Table 3). However, chromatin accessibility was not overlapping with these SNPs in OPCs or MOLs in healthy individuals (Fig. 2b, Supplementary Table 3), which might suggest their accessibility might only increase in MS patients and not in healthy individuals. The outside variant SNP rs6498169^19^, located in between the loci of the master MHC-II regulator *Ciita* and *Socs1*, a major regulator of inflammation, also presented accessibility specifically in OPC- and MOL5/6-EAE, but not in Ctr-OLG or MiGl (Supplementary Fig. 3a, Supplementary Table 3) and the non-MHC SNP rs17493811^18^ within the *Pitpnm2* locus was overlapping mainly with chromatin accessibility in Ctr- and EAE-OPCs. These SNPs might be involved in the modulation of regulatory regions in OLG, leading to altered transcriptional output and ultimately to altered function of OLG in the context of MS.

**Fig. 3.**
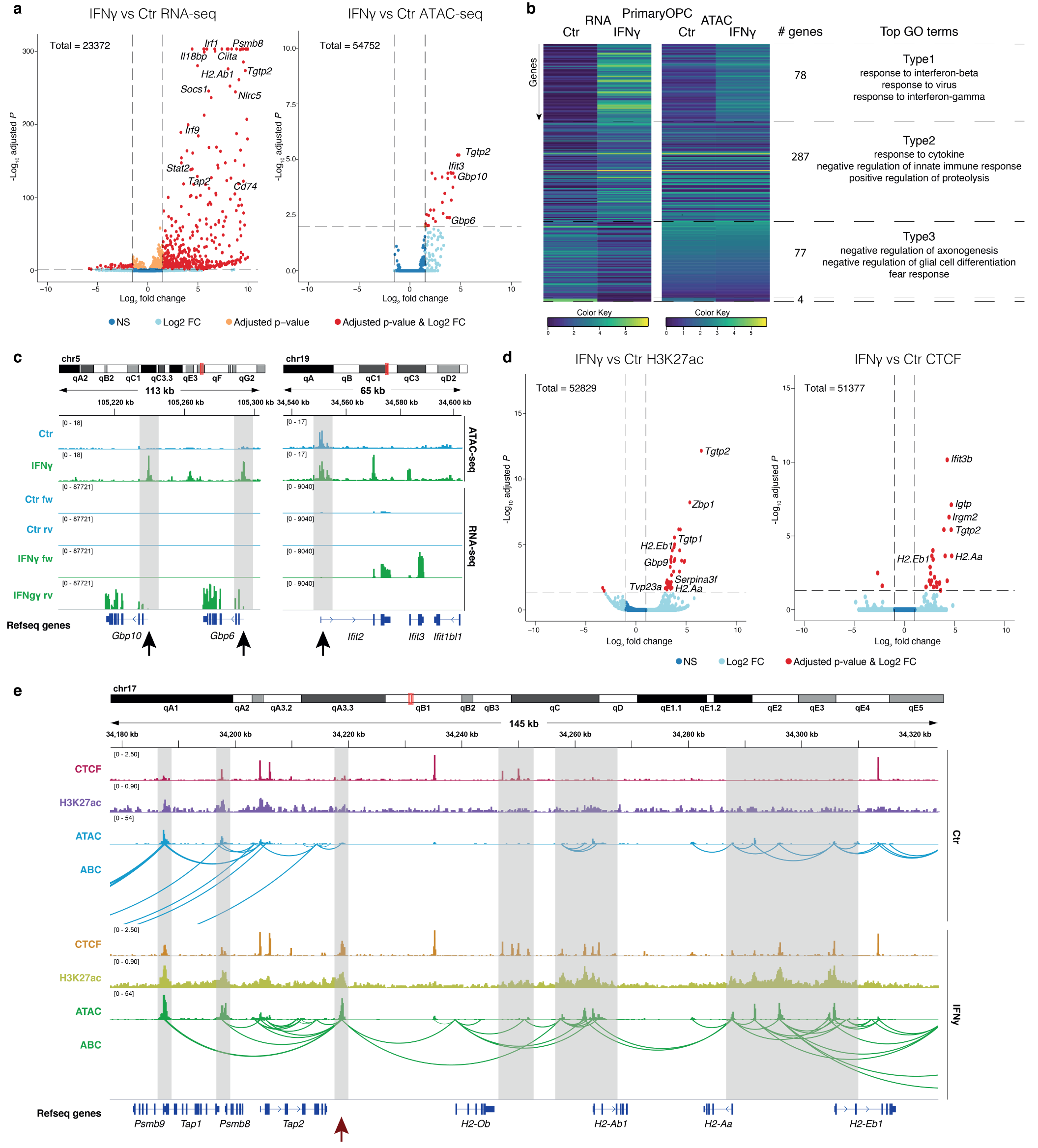
Primed and increased chromatin accessibility at immune gene loci in OPCs upon IFN*γ* treatment. **a**, Volcano plots showing differential gene expression in RNA-seq (left) and chromatin accessibility at promoter regions in ATAC-seq (right) between Ctr-OPCs and OPCs treated with 100 ng/ml IFN*γ* for 48 hours. Genes with statistical significance are shown in orange and genes with statistical significance and log2 fold change above 1.5 are shown in red. **b**, Genes in IFN*γ*-treated and Ctr-OPCs are clustered based on their chromatin activity score (chromatin accessibility over 500 bp promoter region) and gene expression correlation. Top GO terms are shown for Type1 (genes with increased expression and chromatin accessibility upon IFN*γ* treatment), Type2 (genes with increased expression upon IFN*γ* treatment, but no change in chromatin accessibility) and Type3 (genes with reduced expression upon IFN*γ* treatment, but no change in chromatin accessibility). Type4 (genes with reduced expression and chromatin accessibility upon IFN*γ* treatment) had no GO terms because of too few genes in this group. **c**, IGV tracks are shown for ATAC-seq and RNA-seq in IFN*γ*-treated and Ctr-OPCs for selected genes. Highlighted with grey boxes and arrows are regions with differential chromatin accessibility in IFN*γ*-treated OPCs or promoter priming, black arrows indicate promoter regions. Merged tracks of 3 replicates are shown for ATAC-seq and 4 replicates for RNA-seq. **d**, Volcano plots showing differential H3K27ac (left) and CTCF binding (right) between IFN*γ*-treated and Ctr-OPCs, assessed with Cut&Run. 3 replicates were performed. Genes with statistical significance and log2 fold change above 1.5 are shown in red. **e**, IGV tracks showing CTCF binding and H3K27ac occupancy, assessed with Cut&Run, ATAC-seq in IFN*γ*-treated and Ctr-OPCs for MHC-I and MHC-II loci. Predicted enhancer/promoter contacts computed by the activity-by-contact (ABC) model ^16^ are shown. Highlighted with grey boxes are regions with increased H3K27ac, CTCF binding, accessibility and/or predicted interactions in IFN*γ*-treated OPCs. Highlighted with a red arrow is an enhancer region interacting with multiple genes in the MHC-I and MHC-II loci. Merged tracks for 3 replicates per condition are shown.

### Transcription factors involved in immune regulation have increased motif accessibility in EAE-OLG

Transcription factors (TFs) are key modulators of chromatin and transcription by assembling in complexes containing chromatin modifying enzymes and other modulators and recruiting them to specific genomic loci. To explore which TFs might undergo accessibility changes in EAE, we applied chromVAR ^22^ to determine TF motif variability. As expected, we observed clusters of TFs as IRF1, STAT2 and ETS1 with enriched accessible motifs in MiGl, and members of the SOX family, but also ASCL1 and NFIA in Ctr-OPCs and -MOLs (Fig. 2c, Supplementary Table 4). Strikingly, a cohort of TFs presented differential motif activity in OLG in EAE, including TFs as FOS, SMARCC1, and other TFs with known immunoregulatory functions, such as the TF BTB and CNC homology (BACH1), BACH2 and the basic leucine zipper ATF-like TF BATF (Fig. 2c, Supplementary Fig. 3b, c, Supplementary Table 4). We then further analyzed which TF motifs were enriched in EAE compared to CFA-Ctr in OPCs, MOL1/2 and MOL5/6 specifically (Supplementary Fig. 3c, Supplementary Table 4). TFs such as IRF1, STAT2 and KLF4 had enriched motifs in EAE in all three OLG populations and had predicted binding sites in EAE in promoter and enhancer regions of the *Psmb9*-*Tap2* locus (Supplementary Fig. 3c, d). Projection of the TF motif activity on top of the UMAP clustering indicated that TFs as STAT1, STAT3, HIF3a, RELA, SMARCC1 and NFIX presented differential motif accessibility in most of OLG populations in EAE, while other TFs were particularly enriched in specific OLG-EAE populations, such as IRF1 and STAT2 in OPC-EAE, IRF7, FOS, JUNB, KLF13, KLF4, BACH1 and BACH2 in MOL1/2-EAE and SOX8 in all MOL-EAE populations (Fig. 2d, Supplementary Fig. 3b). Most of these TFs exhibited increased expression in OLG from EAE mice, however a subset of them did not, such as SOX8 (in MOL1/2), SMARCC1 (in MOL1/2), NFIX (in OPCs, MOL1/2 and MOL5/6) and BACH1 (in OPCs, MOL1/2 and MOL5/6) (Supplementary Fig. 3c). Thus, TFs as SOX8, expressed throughout the OLG lineage and implicated in OL development ^23^ and MS susceptibility ^18^, might work in concert with TFs other than the ones they usually cooperate with during development and homeostasis, in order to regulate immune gene transcription in OLG in EAE/MS ^1^.

### Activation of primed immune genes in OPCs by IFN*γ*but not dexamethasone

We and others have previously observed that IFN*γ* induces an immune transcriptional program in OPCs ^1,4^, similar to the phenotype observed in the EAE mouse model of MS ^1^. To investigate whether modulation of chromatin accessibility is involved in IFN*γ*-mediated activation of these immune transcriptional programs, we treated primary mouse OPCs with IFN*γ* for 48 hours and performed bulk ATAC-seq and RNA-seq. While *Sox10* (expressed in OLG) and *Ptprz1* (expressed in OPCs) did not change in accessibility or gene expression (Supplementary Fig. 4a), genes with increased expression upon IFN*γ* treatment included interferon response genes *Stat1, Stat2* and *Irf1*, MHC-I and MHC-II genes *H2-k1, H2-q7, H2-ab1, H2-aa*, among others (Fig. 3a, Supplementary Fig. 4b, Supplementary Table 5). The top GO biological terms for genes up-regulated upon IFN*γ*treatment were ‘defense response’, ‘response to cytokine’ and ‘cellular response to cytokine stimulus’ (Supplementary Fig. 4c, Supplementary Table 5). Interestingly, there were fewer chromatin accessibility sites altered upon IFN*γ* treatment, when compared to the number of genes with altered expression (Fig. 3a, Supplementary Table 5). Nevertheless, we did find similar GO-terms when analyzing peaks within proximal (promoter) regions of genes, including ‘response to interferon-beta’, ‘defense response to protozoan’ and ‘response to interferon-alpha’ (Supplementary Fig.4d, Supplementary Table 5). However, we did find more changes in chromatin accessibility at annotated enhancers (Supplementary Fig. 4e, Supplementary Table 5). While not many genes were downregulated upon IFN*γ* (Fig. 3a), the genes that were downregulated were included in GO terms related to system and CNS development (Supplementary Fig. 4c, Supplementary Table 5). Among the genes downregulated were *Sox8, Myrf* and *Plp1*, genes important for OPC specification and differentiation (Supplementary Fig. 4f). These results are consistent with the reported negative impact of IFN*γ* on OPC differentiation ^4,24^. Interestingly, we did not find any loss of accessibility at their promoters or annotated enhancers (Supplementary Fig. 4f).

**Fig. 4.**
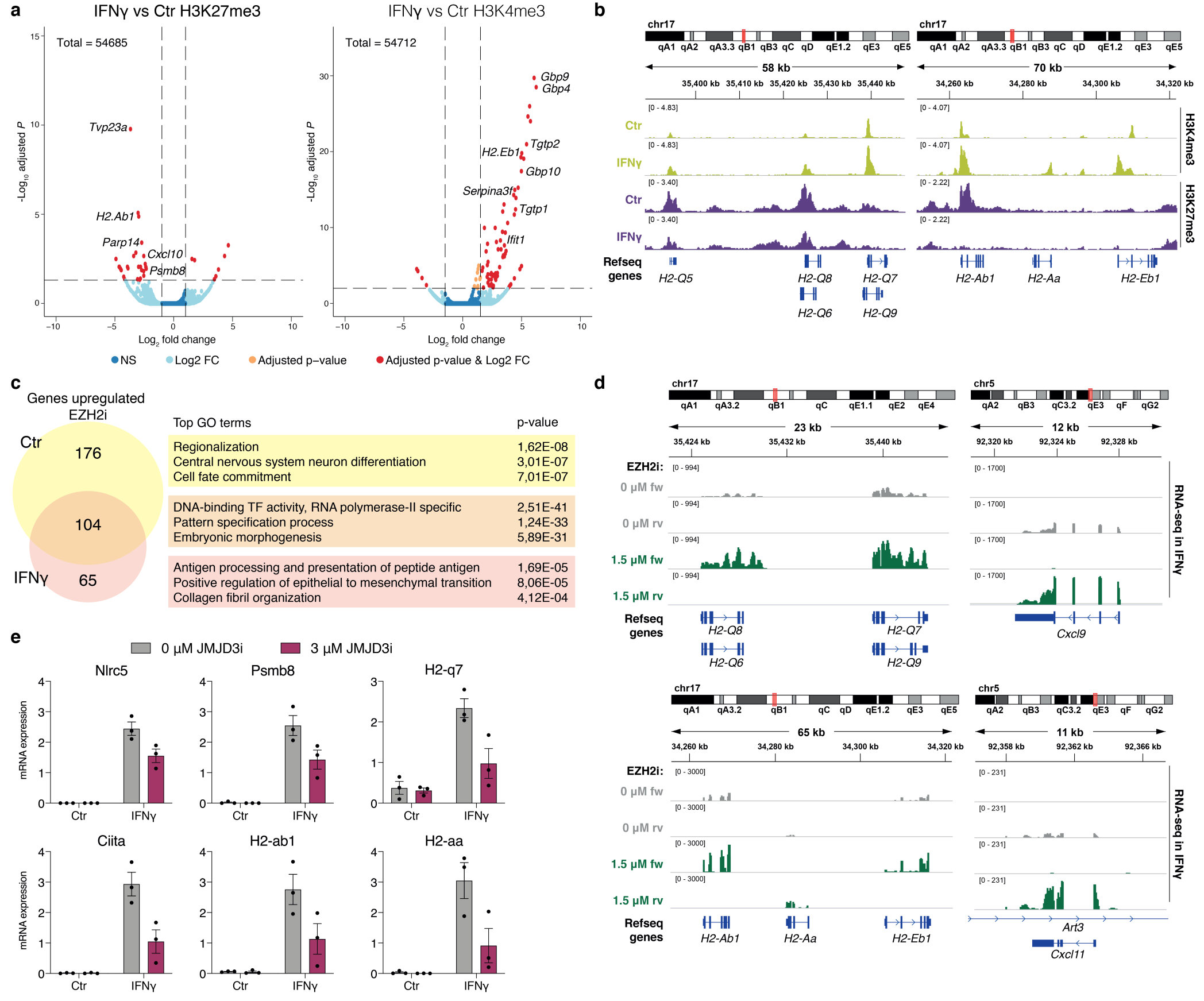
H3K27me3 demethylation is required for IFN*γ* mediated immune gene activation in OPCs. **a**, Volcano plots for H3K27me3 and H3K4me3 in IFN*γ*-treated vs Ctr-OPCs assessed with Cut&Run. Two replicates were performed. Genes with statistical significance are shown in orange and genes with statistical significance and log2 fold change above 1.5 are shown in red. **b**, Cut&Run IGV tracks for MHC-I and MHC-II genes with increased H3K4me3 or decreased H3K27me3 in OPCs upon IFN*γ* treatment. Merged tracks of two replicates are shown. **c**, Venn diagram showing the number of genes enriched in OPCs upon treatment with 1,5 µM EZH2 inhibitor EPZ011989 (EZH2i) for 4 days, with and without subsequent co-treatment with 100 ng/ml IFN*γ* for 6 last hours (and the genes enriched in both) and the top Gene Ontology biological terms for the genes in each category. **d**, RNA-seq IGV tracks for MHC-I, MHC-II and cytokine genes with increased expression upon EZH2i in IFN*γ*-spiked OPCs. Merged tracks of three replicates are shown. **e**, qRT-PCR analysis of MHC-I and MHC-II pathway genes in OPCs upon treatment with 3 µM JMJD3/*Kdm6b* inhibitor GSKJ-4 (JMJD3i) for 2 days, with subsequent co-treatment with 100 ng/ml IFN*γ* for 6 last hours. Error bars represent SEM, three replicates are shown.

Similar to the EAE scATAC-seq data (Fig. 1g), a subset of the upregulated immune response genes had already accessibility in control OPCs at their promoter and only a few genes altered their chromatin accessibility upon IFN*γ*exposure (Fig. 3b). The top GO terms for those type of genes that were already accessible (Type2) were ‘response to cytokine’, ‘negative regulation of innate immune response’ and ‘positive regulation of proteolysis’ (Fig. 3b, Supplementary Table 5). Some of the Type2 genes had enhanced accessibility within their gene body or in the regions upstream of the TSS, pointing towards promoter priming and specific enhancer regulation of expression. Examples of those type of genes are *Irf1, Stat3*, and *Ifit2*, while other genes, such as the *Gbp* gene family, presented increases in both accessibility and gene expression upon IFN*γ* treatment (Fig. 3c). Thus, OPCs already exhibit primed chromatin accessibility in a subset of immune genes prior to inflammatory insults both ex vivo and in vivo.

We had previously observed that treatment of OPCs with the anti-inflammatory corticosteroid dexamethasone (Dexa) led to induction of a subset of genes expressed in specific OL lineage populations in EAE ^1^, involved for instance in cell survival ^25^. Bulk RNA-seq analysis confirmed that many of the genes upregulated upon Dexa treatment were also enriched in the OPC2 and MOL56-EAE-a populations (15 out of 85 genes) ^1^ (Supplementary Fig. 4g-i, Supplementary Table 6). Interestingly, no genes were found downregulated upon Dexa treatment (Supplementary Fig. 4g, h). Top GO biological terms enriched upon Dexa treatment were ‘Regulation of Rho protein signal transduction’, ‘Response to osmotic stress’ and ‘stress fiber assembly’ (Supplementary Fig. 4j, Supplementary Table 6). In contrast to IFN*γ*, we could not find any chromatin accessibility changes upon Dexa treatment (Supplementary Fig. 4h). Thus, IFN*γ* and Dexa regulate different transcriptional networks in OPCs, through distinct chromatin regulatory mechanisms.

### Increased interactions between enhancers and promoters at MHC-I and MHC-II loci in OPCs upon IFN*γ* treatment

A considerable number of immune genes have already promoter accessibility in Ctr OPCs (Fig. 1g, 3b), suggesting that additional mechanisms are required for their transcriptional activation in the context of EAE/IFN*γ* treatment. One of such mechanisms could be the increase of interactions between enhancers and immune gene promoters upon IFN*γ*exposure in OPCs. We therefore profiled H3K27ac with Cut&Run ^26^ to identify active enhancers. An increase of H3K27ac was observed at enhancers and promoters of immune genes, such as *Tgtp1/2, Zbp1, Gbp9, Cd74, Nlrc5, H2-aa* and *H2-eb1* (Fig. 3d, e, Supplementary Table 7). We then used the activity-by-contact model ^16^ to predict promoter-enhancer interactions in both Ctr- and IFN*γ* -OPCs based on chromatin accessibility and H3K27ac. We observed an increase in predicted interactions within and between the two neighboring *Psmb9*-*Tap2* and *H2-ab1*-*H2-eb1* loci (Fig. 3e). An enhancer downstream of *Tap2* harbors a STAT1/STAT2 motif, which has increased chromatin accessibility and H3K27ac upon IFN*γ* treatment, and seems to be the major enhancer connecting the two loci and interacting with most genes in the region (Fig. 3e). The chromatin accessibility (Fig. 1f) and STAT2 motif accessibility (Supplementary Fig. 3d) at this enhancer were also increased in OLG-EAE. CTCF has been recently shown to be required for acute inflammatory responses in macrophages ^27^ and an increase in CTCF mediated promoter-enhancer interactions at the MHC-I and MHC-II loci was previously observed in B-cells ^28,29^. Therefore, we profiled CTCF with Cut&Run upon treatment with IFN*γ* and observed increased binding of CTCF in enhancers and at the promoters of many immune genes (Fig. 3d, e, Supplementary Table 7). These results suggest that altered chromatin accessibility, together with increased CTCF binding, H3K27ac deposition and promoter-enhancer interactions might contribute to the activation of an immune gene transcriptional program in already primed OPCs.

### H3K27me3 demethylation is required for IFN*γ* mediated immune gene activation in OPCs

H3K4me3/H3K27me3 bivalency has been shown to mediate poising of cell-type specific transcription during development ^30^ and immune gene transcription in cancer ^31^. Hence, H3k27me3 mediated repression of poised genes could be another mechanism of immune gene regulation in OPCs. We therefore used Cut&Run to profile H3K27me3 and H3K4me3 in primary OPCs. In Ctr OPCs, H3K27me3 and H3K4me3 bivalency was indeed observed at the promoters of MHC-I and -II genes, including *Tap1, H2-q5, H2-q6, H2-q7, H2-k1* and *H2-ab1* (Supplementary Table 8).Strikingly, upon 48 hours of IFN*γ* treatment, H3K27me3 was reduced at immune gene loci, including *H2-ab1, Cxcl10* and *Psmb8* (Fig. 4a, b, Supplementary Table 7). Reversely, an increase of H3k4me3 is observed at immune genes such as *Gbp4, 9, Tgtp1, 2, Serpina3f, H2-ab1, H2-aa* and *H2-eb1* (Fig. 4a, b, Supplementary Table 7). Thus, resolving bivalency might contribute to the activation of transcription of immune genes in OPCs upon IFN*γ* treatment.

To investigate if H3K27me3 mediated derepression is sufficient for immune gene upregulation we inhibited EZH2, the enzyme responsible for the deposition of H3K27me3 (Supplementary Fig. 5a). Inhibition of EZH2 (EZH2i) with EPZ011989^31^ led to a reduction of H3K27me3 overall (Supplementary Fig. 5b) and at specific genomic loci (Supplementary Fig. 5c, d), and a derepression of TFs involved in specification and morphogenesis (Fig. 4c, Supplementary Fig. 5e, Supplementary Table 9), in line with previous literature showing the importance of EZH2 in the specification o f O PCs ^32^. I mportantly, E ZH2i l ed t o an upregulation of the expression of a subset of immune genes, such as *H2-d1* and *Tap1*, while others remained repressed. Spiking EZH2 inhibited cells with IFN*γ* for 6 hours led to an increased upregulation of MHC-I and MHC-II genes *H2-q6, H2-q7, H2-ab1, H2-aa, H2-eb1* and genes involved in cytokine signaling including *Cxcl9* and *Cxcl11* (Fig. 4c, d, Supplementary Fig. 5e, f, Supplementary Table 9). These results suggest that H3K27me3 removal synergizes with inflammatory c ues a s IFN*γ* t o p romote i mmune gene transcription in OPCs.

Since IFN*γ* treatment of OPCs led to a reduction of H3K27me3 deposition at immune genes, we investigated whether H3K27me3 demethylation is necessary for immune gene upregulation. For this purpose, we used the H3K27me3 specific demethylase inhibitor GSKJ-4, targeting JMJD3 and UTX ^33^. Pre-treatment with GSKJ-4 limited the IFN*γ*-induced upregulation of *H2-q7, H2-ab1* and *H2-aa* (Fig. 4e). MHC-I and MHC-II master regulators *Nlrc5* and *Ciita* showed also decreased upregulation (Fig. 4e). These results show that H3K27me3 removal by JMJD3/UTX upon IFN*γ* exposure is important to incite a full activation of the immune response within OPCs.

## Discussion

OLG have been thought to have a dedicated function in the CNS to produce myelin and allow fast transmission of electrical impulse within the neuronal circuitry. The finding t hat O PCs a nd M OLs c an s tart e xpressing immune genes in the context of MS adds new roles to the functional portfolio of OLG. OPCs with immunological properties can activate CD4 T-cells, inducing their proliferation, survival and production of cytokines ^1^, and CD8 T-cells, triggering a negative feedback loop that can lead to OPC cell death ^4^. Interestingly, this capacity to activate an immune transcriptional program might not be circumscribed to MS, but also occurs in Alzheimer’s disease ^34^ and in aging ^35–38^.

In this study, we find t hat O LG a re a lready p rimed at the chromatin level, in order to be able to rapidly activate immune gene programs in the context of disease. This is reflected both on the open chromatin state in several immune gene loci and the bivalency for H3K4me3 and H3K27me3. The transition to an immune active state involves further chromatin accessibility in additional gene loci and resolution of the bivalency. Our results suggest that IFN*γ* plays an important role in the transition of OLG into immune states, by leading to a rearrangement of CTCF binding and remodeling of promoter-enhancer interactions. While our data supports the role of distal enhancers in establishing cell identities ^9,10^, it also highlights the importance of promoters not only in this process, but more importantly, in the transition between functional and disease-specific c ell s tates w ithin t he same cell identity.

Single nucleotide polymorphisms (SNPs) in MS are in most cases located nearby genes involved in immune regulation ^18^. As such, susceptibility for MS has been mainly linked to immune cells within the CNS or in the periphery ^18^. Our data indicate that a subset of SNPs present in MS patients are located in regulatory regions that present open chromatin in OLG in homeostatic conditions in mouse and in healthy individuals in human, or that exhibit increased accessibility in OLG in the EAE mouse model of MS. Some of the genes that are associated with these regulatory regions also have increased expression in EAE ^1^. Recent findings also suggest that SNPs located in regulatory regions of genes involved in transcriptional elongation might be involved in dysregulation of OL differentiation in the context of MS ^19^. Accordingly, susceptibility for MS might lead to disease onset, progression or remission by the activation of abnormal immune and non-immune transcriptional programs not only in immune cells but also in OLG, which therefore constitute novel targets for immunological-based therapies for MS.

IFN*γ* is a key inducer of immune states in OLG ^1,2,4,5^. Here we found that one of the mechanisms by which IFN*γ*promotes such cellular state transitions is by inducing demethylation of H3K27me3 at immune gene promoters in OPCs. We also observed that this process can be antagonized by an inhibitor of JMJD3 and UTX, GSKJ-4. This inhibitor has been previously shown to be a clinically relevant target for diffuse intrinsic pontine glioma ^39^ and T-cell acute lymphoblastic leukaemia ^40^. As such, compounds as GSKJ-4 might constitute important tools for therapeutic efforts targeting the response of OLG to inflammatory signals in the context of MS.

## ACKNOWLEDGEMENTS

We would like to thank Tony Jimenez-Beristain and Marek Bartosovic for assistance in experiments, Jaromir Mikes and the Biomedicum FACS facility for support in FACS sorting, the Single Cell Genomics Facility, the staff at Comparative Medicine-Biomedicum and Sten Linnarsson’s lab for additional support. The authors acknowledge support from the National Genomics Infrastructure in Stockholm funded by Science for Life Laboratory, the Knut and Alice Wallenberg Foundation and the Swedish Research Council, and Swedish National Infrastructure for Computing /Uppsala Multidisciplinary Center for Advanced Computational Science for assistance with massively parallel sequencing and access to the UPPMAX computational infrastructure. A.M.F. is supported by the European Committee for Treatment and Research in Multiple Sclerosis (ECTRIMS) and a fellowship from “La Caixa” Foundation (ID 100010434) with the reference LCF/BQ/PI19/11690005. E.A. is funded by the European Union, Horizon 2020, Marie-Sklodowska Curie Actions, and grant SOLO, number 794689. Work in G.C.-B.’s research group was supported by the Swedish Research Council (grant 2015-03558 and 2019-01360), the European Union (FP7/Marie Curie Integration Grant EPIOPC, Horizon 2020 Research and Innovation Programme/European Research Council Consolidator Grant EPIS-cOPE, grant agreement number 681893), the Swedish Brain Foundation (FO2017-0075), the Ming Wai Lau Centre for Reparative Medicine, the Swedish Cancer Society (Cancerfonden; CAN2016/555), Knut and Alice Wallenberg Foundation (grant 2019-0107), The Swedish Society for Medical Research (SSMF, grant JUB2019), Ming Wai Lau Centre for Reparative Medicine and Karolinska Institutet. H.Y.C. is supported by NIH RM1-HG007735 and is an Investigator of the Howard Hughes Medical Institute. X.C is supported by the Swedish Research Council (VR-2017-02074). M.R.C is supported by the National Institute on Aging (K99AG059918). T.J.M. is supported by NIH P50 NS062684. This manuscript was formatted using the BioRxiv template in the Overleaf software (Author: Ricardo Henriques, License: Creative Commons CC BY 4.0).

## AUTHOR CONTRIBUTIONS

G.C-B, M.M. and E.A. conceived and designed the project, interpreted data and wrote the manuscript. M.M, M.K., C.A.v.T., A.H, and A.M.F performed experiments. E.A. performed computational analyses, M.K. performed ABC analysis. M.R.C, T.J.M. and H.Y.C. performed and provided human scATAC-seq data. X.C. and H.Y.C. provided valuable advice and input for Pi-ATAC experiments. G.C-B supervised the project. All the authors critically reviewed the manuscript and approved the final version.

## DATA AVAILABILITY

The scATAC-seq dataset can be explored at the following web resource: https://castelobranco.shinyapps.io/SCATAC10X_2020/. Raw data is currently being deposited in GEO (GSE154175) and will be made available at a later stage.

## DECLARATION OF INTERESTS

H.Y.C. is a co-founder of Accent Therapeutics, Boundless Bio, and an advisor to 10x Genomics, Arsenal Biosciences, and Spring Discovery. The other authors declare no competing interests.

## Methods

### Animals

The mouse line used in this study was generated by crossing Sox10:Cre animals ^7^ (The Jackson Laboratory mouse strain 025807) on a C57BL/6j genetic background with RCE:loxP (EGFP) animals ^8^ (The Jackson Laboratory mouse strain 32037-JAX) on a C57BL/6xCD1 mixed genetic background. Females with a hemizygous Cre allele were mated with males lacking the Cre allele, while the reporter allele was kept in hemizygosity or homozygosity in both females and males. In the resulting Sox10:Cre-RCE:LoxP (EGFP) animals the entire OL lineage was labeled with EGFP. Breeding with males containing a hemizygous Cre allele in combination with the reporter allele to non-Cre carrier females resulted in offspring where all cells were labeled with EGFP and was therefore avoided.

For primary cell culture, animals of both sexes were sacrificed at P4-P6. For EAE experiments, males between 9 and 13 weeks old were used for immunization and sacrificed 10-17 days after, at the peak of the disease. Mice were housed to a maximum number of 5 per cage in individually ventilated cages with the following light/dark cycle: dawn 6:00-7:00, daylight 7:00-18:00, dusk 18:00-19:00, night 19:00-6:00. All experimental procedures on animals were performed following the European directive 2010/63/EU, local Swedish directive L150/SJVFS/2019:9, Saknr L150 and Karolinska Institutet complementary guidelines for procurement and use of laboratory animals, Dnr. 1937/03-640. The procedures described were approved by the local committee for ethical experiments on laboratory animals in Sweden (Stockholms Norra Djurförsöksetiska nämnd), lic. nr. 131/15 and 144/16.

### Experimental Autoimmune Encephalomyelitis (EAE)

For the induction of EAE, the mouse model of MS, animals were injected subcutaneously with an emulsion of MOG35-55 peptide in complete Freud’s adjuvant (CFA; EK-2110 kit from Hooke Laboratories) followed by intraperitoneal injection with pertussis toxin in PBS (200 ng per animal) on the day of immunization and with 24 hours delay (according to manufacturer’s instructions). Control animals underwent the same treatment, but CFA without MOG35-55 peptide (CK-2110 kit from Hooke Laboratories) was used instead. Spinal cord and brains were collected at the peak of the disease when clinical score 3 (representing limp tail and complete paralysis of hind legs) has been reached. Animals that did not reach this clinical score were not analyzed in this study.

### Tissue dissociation for single-cell ATAC-seq experiment

Mice were sacrificed with a ketaminol/xylazine intraperitoneal injection followed by intracardiac perfusion with PBS. Brain and spinal cords were collected and dissociated using the adult brain dissociation kit (130-107-677, Miltenyi), following manufacturer’s instructions which included myelin debris removal but not the red blood cell removal step.

### Single-cell ATAC-seq (10x Genomics)

Immediately after dissociation, cells were stained with DAPI (0.5 µg/ml, D9542, Sigma) and sorted on a FACS Aria III cell sorter (BD Biosciences). Sox10-GFP+/DAPI-cells were collected in PBS + 0.5% BSA and pooled with Sox10-GFP-/DAPI-cells with a 4:1 ratio. The pool of cells was then lysed and washed according to the Demonstrated Protocol: Nuclei Isolation for Single cell ATAC Sequencing (10x Genomics) as follows: the cells were centrifuged for 10 minutes at 300xg and 4°C, resuspended in ATAC lysis buffer (containing 0.1% IGEPAL (CA-630), 0.1%Tween-20, 0.01% Digitonin, 1% BSA, 10 mM Tris-HCl pH 7.4, 10 mM NaCl, 3 mM MgCl2) and incubated on ice for 3 minutes. After the incubation, wash buffer (containing 0.1%Tween-20, 1% BSA, 10 mM Tris-HCl pH 7.4, 10 mM NaCl, 3 mM MgCl2) was added on top without mixing and the nuclei were centrifuged for 5 min at 500xg and 4°C. Nuclei were washed once in Diluted Nuclei buffer (10x Genomics) containing 1% BSA, and incubated for 60 minutes at 37°C in tagmentation mix (10x Genomics). The Chromium Single Cell ATAC v1 Chemistry was used to create single-cell ATAC libraries. Two EAE and four CFA-Ctr animals were used for independent replicates. Libraries were sequenced on an Illumina Novaseq 6000 with a 50-8-16-49 read set-up and a minimum of 25 000 read pairs per cell.

### Single-cell ATAC-seq (Pi-ATAC ^12^)

Immediately after dissociation, cells were fixed in 1%formaldehyde (28906, ThermoFisher Scientific) for 10 minutes and quenched with glycine (125 mM) for 5 minutes at room temperature and then washed and stored in 0.5%BSA in PBS with 0.1% sodium azide at 4°C until further processing. The cells were counted and aliquots of 500.000 cells were centrifuged for 10 minutes at 1000xg and room temperature. Cells were resuspended in lysis buffer (containing 0.05% IGEPAL (CA-630), 10 mM Tris-HCl pH 7.4, 10 mM NaCl, 3 mM MgCl2) and incubated for 5 minutes at room temperature. After a 20-minute centrifugation at 1000xg at room temperature, the cells were incubated with anti-GFP antibody (FITC conjugated, 1:100, Ab6662, Abcam) and DAPI (0.5 µg/ml) in PBS containing 5% BSA for 20 minutes at room temperature. The cells were centrifuged for 10 minutes at 1000xg and resuspended in tagmentation mix (dH2O, 2x TD buffer ^41^ and Tn5 enzyme ^42^) and incubated for 30 minutes at 37°C. For both lysis and tagmentation buffers the volume was scaled up to match the number of cells (50 µl per 50.000 cells). Tagmentation reaction was stopped by addition of 40 mM EDTA. Cells were then centrifuged for 10 minutes at 1000xg and room temperature and resuspended in PBS + 0.5% BSA. Sox10 GFP+/DAPI+ cells were sorted on an Influx (BD Biosciences) in reverse crosslinking buffer ^12^ with single cells in each well of 96 well plates. Also Sox10 GFP-/DAPI+ cells were sorted as a negative control for the OL lineage. For reverse cross-linking, the plates were incubated overnight at 65°C ending with 10 minutes at 80°C the next day. PCR master mixes (NEBNext High fidelity, M0541S, NEB) containing unique barcoding primers per well were dispensed on top of the reverse crosslinking buffer and DNA was amplified with the following cycling conditions: 72°C for 5 min, 98°C for 30s; 20 cycles at 98°C for 10s, 63°C for 30s and 72°C for 1 min. PCR products were purified with the MinElute purification kit (Qiagen) and then PAGE purified to remove adapter dimers. Three EAE and two CFA-Ctr animals were used for independent replicates. Libraries were sequenced on an Illumina Hiseq 2500 with a 50-8-8-50 read set-up and a minimum of 25 000 read pairs per cell.

### Tissue dissociation for primary OPC cultures

Brains from P4-P6 mouse pups were collected and dissociated with the neural tissue dissociation kit (P; 130-092-628, Miltenyi) according to manufacturer’s protocol. OPCs were obtained by either FACS with Sox10-GFP+ selection or by MACS with CD140a microbeads (Cd140a microbead kit, 130-101-547, Miltenyi). For each experiment, multiple brains were pooled to obtain a sufficient number of cells.

### Primary OPC culture

Cells were seeded on poly-L-lysine (P4707, Sigma) coated plates (150 000 cell in 3.8cm^2^ -12 well) and grown with OPC proliferation media consisting of DMEM/F12, Gluta-MAX (10565018, ThermoFisher Scientific), N2 supplement (17502001, ThermoFisher Scientific) NeuroBrew 21 (130-097-263, Miltenyi), penicillin-streptomycin (50 Units/ml penicillin, 50 µg/ml streptomycin, 15140122, ThermoFisher Scientific), PDGFaa (20 ng/ml, 315-17, PeproTech) and bFGF (40 ng/ml, 100-18B, PeproTech). Cells were treated with either dexamethasone (1 µM, D4902, Sigma) or IFN*γ*(100 ng/ml, 485-MI-100, R&D) for 48 hours.

### EZH2 inhibition

OPC primary cells were treated with EZH2 inhibitor (EPZ011989, 1.5 µM, S7805, Selleckchem) for 4 days, renewing the inhibitor every 48 hours. After 4 days, IFN*γ*(100 ng/ml) was added for 6 hours together with 1.5 µM EZH2 inhibitor.

### JMJD3 inhibition

OPC primary cells were treated with JMJD3 inhibitor (GSKJ-4, 3 µM, SML0701, Sigma) for 48 hours, adding IFN*γ* (100 ng/ml) for the last 6 hours together with 3 µM GSKJ-4 inhibitor.

### Bulk ATAC-seq

ATAC-seq was performed as previously described ^43^ with minor adaptations. Primary OPCs were incubated with TrypLE (Gibco 12605010) at 37°C for 5 minutes and collected in cell culture media. 60 000 cells per condition were washed with PBS and lysed with lysis buffer (containing 0.1%IGEPAL (CA-630), 10 mM Tris-HCl pH 7.4, 10 mM NaCl, 3 mM MgCl2) and centrifuged for 20 minutes at 500xg and 4°C. Cells were then resuspended in tagmentation mix (dH2O, 2x TD buffer ^41^ and Tn5 enzyme ^42^) for 30 minutes at 37°C. The DNA was purified pre- and post-PCR with the MinElute purification kit (Qiagen) and then PAGE purified to remove adapter dimers. Three replicates per condition were performed with primary OPCs obtained from different litters. Libraries were sequenced on an Illumina Novaseq 6000 with a 50-8-8-50 read set-up.

### RNA extraction, cDNA synthesis and qRT-PCR

Cells were collected with qiazol (Qiagen) and stored at -80°C until further processing. RNA was extracted with the miRNeasy mini kit (Qiagen) according to manufacturer’s instructions. Contaminating DNA was degraded by treatment of the samples with RNase-free DNase (Qiagen) in column. 350 ng RNA was used to synthesize cDNA with the High-Capacity cDNA Reverse Transcription kit (Applied Biosystems) including RNase inhibitor (Applied Biosystems), with annealing for 10 minutes at 25°C and extending for 2 hours at 37°C and inactivation for 5 minutes at 85°C. The cDNA was diluted 1:5 in H2O and 2,5 µl was used in the qRT-PCR reactions with 5 µl of Fast SYBR® Green Master Mix (Applied Biosystems) and 5 pmol of each primer in a final volume of 10 µl. The reactions were run on a StepOnePlus™ System (Applied Biosystems) in duplicate and with reverse transcriptase negative reactions to control for genomic DNA. The running conditions were 20 seconds at 95°C, followed by 40 cycles of 3 seconds of 95°C and 30 seconds of 60°C, then 15 seconds at 95°C, 1 minute at 60°C and 15 seconds at 95°C. Melt curves were generated to control for primer dimers and gene-specific peaks. Relative standard curves for each gene were generated to obtain relative expression values.*Ubc* and *b-Act* were run as housekeeping genes. Expression levels were then calculated by dividing the relative expression value by the value of the geometric mean of the housekeeping genes. Samples were normalized per experiment. Three replicates per experiment were performed with primary OPCs obtained from different litters. Primer sequences used are listed in Table 1.

### RNA-seq

0.1-1 µg of RNA was used to make RNA-seq libraries with the TruSeq Stranded Total RNA Library Prep Gold kit (Illumina) according to manufacturer’s instructions. Four replicates per condition for the IFN*γ*/Dexa experiment and three replicates for the EZH2i experiment were performed with primary OPCs obtained from different litters. Libraries were sequenced on an Illumina Novaseq 6000 with a 150-8-8-150 read set-up.

### Cut&Run

Cut&Run was performed as previously described ^26^ with minor adaptations.Primary OPCs were incubated with TrypLE (Gibco 12605010) at 37°C for 5 minutes and collected in cell culture media. Cells were centrifuged for 5 minutes at 300xg and room temperature and then resuspended in wash buffer (20 mM HEPES pH 7.5, 150 mM NaCl, 0.5 mM Spermidine, 0.01% BSA, 1x Roche Complete Protease Inhibitor tablet). 250 000 cells per condition were centrifuged for 3 minutes at 600xg and room temperature and resuspended in wash buffer. Activated Concanavalin-A beads (Bangs Laboratories BP531) in binding buffer (20 mM HEPES pH 7.5, 10 mM KCl, 1 mM CaCl2, 1 mM MnCl2) was added to each condition and incubated for 10 minutes on a rotator at room temperature. Beads with nuclei were now kept on a magnetic stand and washed with Dig-wash buffer (0.05% digitonin in wash buffer). After discarding the liquid, beads were resuspended with primary antibody in antibody buffer (2 mM EDTA in Dig-wash buffer) and incubated overnight at 4°C on a nutator. Then beads were washed with freshly prepared Dig-wash buffer and incubated with 2 µg/ml Protein A-MNase ^44^ in Dig-wash buffer for 1 hour at 4°C on a nutator. After two washes with Dig-wash buffer and one wash with low-salt rinse buffer (20 mM HEPES pH 7.5, 0.5 mM spermidine, 0.05% digitonin) ice-cold incubation buffer (3.5 mM HEPES pH 7.5, 10 mM CaCl2, 0.05% digitonin) was added to the beads which were subsequently placed in a metal block in an ice-water bath maintained at 0°C for 5 minutes. The beads were then placed on a magnet stand and liquid discarded. Stop buffer (170 mM NaCl, 20 mM EGTA, 0.05% digitonin, 50 µg/mL RNase A, 25 µg/mL glycogen, 2 pg/ml Yeast spike-in DNA) was added to the beads and incubated for 30 minutes at 37°C. Beads were placed on a magnet stand and supernatant was collected. 2 µL of 10% SDS and 2.5 µL of 20 mg/ml Proteinase K was added to the supernatant and incubated for 1 hour at 50°C. DNA from the samples was purified using the MinElute PCR purification kit (Qiagen) according to manufacturer’s instructions. Antibodies were used against H3K27me3 (Cell Signaling 9733S; rabbit; 1 µg), H3K4me3 (Diagenode C15410003-50; rabbit; 1 µg), H3K27ac (Abcam ab177178; rabbit; 1 µg) and CTCF (Cell Signaling 3418S; rabbit; 1:100).

Sequencing libraries were prepared using KAPA HyperPrep kit (Roche 07962363001) and KAPA Unique Dual-Indexed Adapters (Roche 08861919702) according to manufacturer’s instructions, but with the following adjustments: two post-adapter ligation clean-ups were performed using 0.7x and 1.1x AMPure XP beads respectively. PAGE purification was performed on the post-amplification libraries to remove remaining adapter dimers. Two or three replicates per condition for the IFN*γ* experiment and three replicates for the EZH2i experiment were performed with primary OPCs obtained from different litters. Libraries were sequenced on the Illumina Novaseq 6000 with a 50-8-8-50 read setup.

### Western Blot

Cells were collected in 2x Laemmli buffer (120 mM Tris-HCl pH 6.8, 4% SDS, 20% glycerol) and sonicated for 5 minutes at high power with 30s on/off cycles at 4°C. Protein concentrations were measured with nanodrop and equalized with 2x Laemmli buffer.Bromophenol blue (0.1%) and B-Mercaptoethanol (10%) were added to the protein prior to a 5-minute incubation at 95°C to denature the protein. Equal volumes were loaded on 4-20% Mini-Protean TGX precast protein gels (4561094, Bio-Rad) and transferred to a PVDF membrane (GE Healthcare) activated in methanol. Membranes were then blocked in blocking buffer (containing TBS, 0.1% Tween-20 and 5% BSA) for 1 hour at room temperature and incubated overnight with primary antibody (diluted in blocking buffer) at 4°C. The membranes were then washed 3 times 10 minutes in TBS-t (TBS, 0.1% Tween-20) and incubated with a horseradish peroxidase-conjugated secondary antibody for 2 hours at room temperature. Proteins were exposed with ECL prime (GE Healthcare) at a ChemiDox XRS imaging system (Bio-Rad).Primary antibodies were used against H3K27me3 (rabbit monoclonal, 9733S, Cell Signaling, 1:1000) or GAPDH (rabbit monoclonal, 5174S, Cell Signaling, 1:1000) and as secondary antibody anti-rabbit (A6667, Sigma, 1:5000).

## Computational methods

### scATAC-seq 10X Genomics preprocessing

scATAC-seq (10X Genomics) data was processed with default parameters with cellranger-atac (version 1.2.0) *count* function.Reads were aligned to mm10 reference genome. As part of cellranger-atac pipeline, peaks were called individually for each of the samples and then merged. Normalized single peak-barcode matrix combining all the samples was calculated with the parameter cellranger-atac *aggr –normalize=depth*, by subsampling all the fragments to the same effective depth to avoid batch effects introduced by sequencing depth, which resulted in a median fragments per cell of 21836. The number of fragments in peaks, the fraction of fragments in peaks and the ratio of reads in ENCODE blacklist sites computed by cellranger-atac were used as QC metrics in downstream processing with the package Signac v0.25 https://satijalab.org/signac/. scATAC 10X Genomics cells with the following metrics were selected; peak_region_fragments > 1000 & peak_region_fragments < 20000 & pct_reads_in_peaks > 15 & blacklist_ratio < 0.05 & nucleosome_signal < 10 & TSS_enrichment_score > 2, which resulted in 4895 cells.

### scATAC-seq Pi-ATAC preprocessing

scATAC-seq (Pi-ATAC) data was processed following the https://carldeboer.github.io/brockman.html_pipeline ^45^. Reads were trimmed and aligned to mm10 reference genome using Bowtie2^46^. Reads with alignment quality less than Q30, incorrectly paired and mapped to mitochondria were discarded.Duplicates were removed using Picard tools. Peaks were called individually for each sample using MACS2^47^(https://github.com/macs3-project/MACS)with the following parameters *-q 0.05 –nomodel -molambda –shift -100 –extsize 200 -call-summits*. Called peaks were merged using bedtools *mergebed* and the peaks overlapping ENCODE blacklisted regions were removed. The summit peaks were resized and extended to the same size and used as input in chromVAR ^22^ to get the fragment counts in peaks, using as input all the individual cell bam files. Fraction of fragments per peak was calculated and filtered using chromVAR resulting in 1029 cells. The output fragments matrix and peak annotations were used as input for Seurat ^11^ to perform non-linear dimension reduction, normalization and clustering.

### Normalization and clustering of scATAC-seq

Normalization and linear dimensional reduction were performed with Signac ^11^.Signac first performs a term frequency-inverse document frequency normalization (TF-IDF), which normalizes across cells and peaks. Then, a feature selection was performed using all the peaks as input. The dimensional reduction was performed on the TF-IDF normalized matrix with the selected peaks using a singular value decomposition (SVD). RunUMAP, FindNeighbors and Findclusters functions from Seurat ^11^ were used for clustering and visualization with 30 dimensions.

### Gene activity scores and integration with scRNA-seq data

We assumed correlation between promoter accessibility and gene expression. First, different promoter lengths were tested (2Kb, 1Kb, 500bp and including the region around the TSS). We extracted the gene coordinates and extended them to include the different promoter lengths. Then, the number of reads from the pooled scATAC-seq samples intersecting the coordinates were counted to calculate a pseudobulk accessibility score. Using scRNA-seq from Falcao *et al*. ^1^, we generated pseudo-bulk scRNA-seq signal for each of the annotated OLG and MiGl cell types and calculated normalized gene expression for the pooled cells. We directly correlated the pooled scATAC accesibility score over the tested promoter regions and pooled gene expressions. The promoter length with highest Spearman correlation coefficient between pseudobulk scATAC and scRNA samples was selected, 500bp. However, 500bp promoters only showed a slight improvement in the correlation value, 500bp Spearman correlation rho for 500bp ∼0.62, 1Kb ∼0.59 and 2Kb ∼0.56.

Final gene activities were computed over the 500bp region upstream the TSS of annotations with the ENSEMBL79 biotype protein_coding using Signac. scATAC-seq cells were annotated based on scRNA-seq data from Falcao *et al*. ^1^ Shared correlation patterns between gene activity and the scRNA-seq annotated expression matrix from Falcao *et al*. ^1^ in Signac/Seurat ^11^ with FindtTransferAnchors refer-ence precomputed scRNA-seq and query scATAC activity scores. Using the classification score of minimum 0.4 the scATAC-seq cells were annotated. Manual checking of the classification scores identified mismatches from incorrect classified cells, where some cells from EAE or Ctr were incorrectly classified, we manually curated the final annotations and discarded cells that showed ambiguity. Single cell tracks were obtained with samtools 1.10 using the CB tag from cellranger-atac aligned bam files and the cluster cell type annotations.

### Differentially accessible peaks

Differential accessibility was calculated between cell type clusters and within cell types between EAE and Ctr conditions with Signac with parameters *min.pct = 0.2, test.use = LR, latent.vars = peak_region_fragments*. For each of the peaks the closest gene was found using closestFeature function combined with EnsDb.Musmusculus.v79 annotations. Identified genes were used in downstream analyses as for instance Gene Ontology. Unique peaks and gene lists per cluster cell type were obtain by comparing unique lists of candidates with adj-pval less than 0.05.

### Gene ontology analysis

GO analysis was performed with ClueGO (version 2.5.5) ^48^ plug-in Cytoscape (version 3.7.2) ^49^ with the following settings: GO Biological process, minimum GO level: 3, Max GO level: 8, minimum number of genes: 3, minimum percentage 4.0, correction methods: bonferroni step down, pvalue cutoff: 0.05.

### CorrelationbetweenscRNA-seqandscATAC-seq heatmaps

The pseudobulk scATAC reads per annotated cell type were intersected with Bedtools bedcoverage with 500bp promoter regions mm10 reference genome EnsEmbl v75 and normalized by total number of reads and region length to get a normalized activity score comparable to normalized expression values. Pseudobulk gene expression and pseudobulk activities at promoters were combined and clustered based on enrichment in EAE compared to Ctr. 4 types of genes were defined based on the expression and activity values, the non-redundant list of genes for each type was used to do gene ontology analysis.

### scATAC peaks annotations

Peak were annotated using HOMER v4.11^50^ with annotatePeaks.pl and gencode.vM20 annotations for all the peaks used in the analysis and for the set of peaks that showed differential accessibility between EAE- and Ctr-OLG. For the donut plots the basic annotations are shown.

### MS associated SNPs

To enable comparison between mouse open chromatin regions and human MS associated SNPs, liftOver was used with parameters *minMatch=0.5* to convert mm10 coordinates to hg19 genomic coordinates. Then, as a double check, we reciprocal lifted back the coordinates to mm10 and retrieved only the peaks that mapped to the original position.In order to define the set of peaks used in the analysis, the properly aligned reads from annotated OLG cell types and MiGl were combined to generate pseudobulk ATAC-seq bam files specific for each cell type. Then, peak calling was performed for each of the annotated cell types, MACS2 with parameters *-q 0.05 –nomodel -molambda –shift -100 –extsize 200*.Peaks were sorted and merged to non-overlapping meta peaks. Using bedtools *intersect -wo* the set of SNPs overlapping open chromatin regions were retrieved.On Figure 2a, the signal of the scATAC-seq is shown on its original mm10 reference and the SNP coordinates on their original hg19 reference. Oligodendrocyte, microglia and OPC scATAC-seq peaks from healthy donors in hg38^17^ were liftOver to hg19.Using bedtools *intersect –wo* with the selected SNPs with evidence from mouse scATAC-seq liftOver peaks the intersecting hg19 peaks were retrieved.

Celltypespecific(CTS)LDscoreregression (https://github.com/bulik/ldsc/wiki/Cell-type-specific-analyses) ^20^ was used to estimate the enrichment coefficient of each OLG and MiGl cluster with different sets of Ctr peaks and a merged set of all significant peaks in the study as background.Gene set enrichments were calculated with MAGMA ^21^ v1.06 with the following parameters; –annotate_window = 10,1.5 –gene_loc = NCBI37.gene.loc with specifically formatted summary statistics.Summary statistics and SNPs locations were obtained from ^18^ and ^19^.

### TF motifs differential accessibility

Motif activity between single cells was calculated using chromVAR ^22^. Motif variability was calculated for all cells on the selected peaks from Seurat/Signac ^11^ analysis with chromVARmotifs library, mouse_pwms_v2, motifs.For visualization purposes the top 100 most variable motifs were selected to build a matrix of the normalized deviation scores, zscores, as is shown in the heatmap in Figure 2b. Deviation zscores from chromVAR are shown on the UMAP coordinates Figure 2c.

Signac was used to identify the overrepresented TF motifs in the sets of differentially accessible peaks between cell types and between EAE vs. Ctr in OLG. Signac performs a hypergeometric test to get the probability of observing a specific motif at a given frequency by chance. The motif enrichment was performed with the chromVARmotifs PWM mouse_pwms_v2 library. Then, we cross checked expression levels of the significantly enriched motifs in the scRNA-seq data from ^1^ to select a set of TFs for further analysis.

For each of the selected TFs the PWM meme files were retrieved from MEME database. Then, FIMO ^51^ was run with mm10 genome reference and the output gff file were converted to bed files with all the genome wide motifs coordinates. In order to predict the binding sites specific to EAE and Ctr OLG, Centipede ^52^ was run using the pseudobulk ATAC signal from each of the OLG clusters on the genome-wide motifs coordinates. Predicted binding sites were selected with a probability score higher than 0.99.

The distribution of the binding sites was annotated using HOMER ^50^ annotatepeaks.pl with the basic annotations. The closest gene were assigned to the predicted binding sites to count the number of genes per cell type cluster for each specific TF.

### Bulk ATAC-seq alignment and peak calling

ATAC-seq samples were processed separately following the standard ENCODE pipeline for ATAC-seq samples, https://www.encodeproject.org/pipelines/ENCPL792NWO/. Adapters were detected and trimmed and reads were alignedto mousegenome(GRCm38/mm10)using Bowtie2, with default parameters.After filtering mitochondrial DNA, reads properly paired were retained and multimapped reads,with MAPQ<30,were removed using SAMtools ^53^.PCR duplicates were removed using MarkDuplicates (Picard,latest version 1.126), http://broadinstitute.github.io/picard/. ATAC-seq peaks were called using MACS2 https://github.com/taoliu/MACS, with parameters *-g mm -q 0.05-nomodel-shift -100-extsize 200 -B -broad*. For visualization and further analyses, the replicates were merged using SAMtools ^54^ and tracks were normalized using deeptools bamcoverage.

### Bulk RNA-seq alignment and differential gene expression

The bulk RNA-seq samples were preprocessed for adapter/quality trimming and then aligned to the transcriptome using STAR ^55^ version 2.7 –quantMode –sjdbOverhang 99 with EnsEMBLv75 gtf annotations.Only uniquely mapped reads were retained for downstream analysis using SAMtools. Aligned samples were converted to bedgraph files using Deeptools bamcoverage for each strand and normalized to total of reads. Filtered fastq files were used in Salmon 0.8.2 to recover the raw reads counts and transcript per million (TPM) values per transcript and gene.The differential gene expression analysis was performed with Deseq2^54^. Results from differential expression were plot using Enhancedvolcano package with log2 fold change and adjusted p-val from Deseq2.

### CUT&RUN alignment and processing

Cut&Run samples were processed with the pipeline CUT&RUNTools that includes reads trimming, alignment (Bowtie2mm10reference genome) and peak calling with MACS2 https://bitbucket.org/qzhudfci/cutruntools/src/master/. For TF Cut&Run samples the narrow peaks with <120bp fragments were used and for histone modifications broad peaks from all the fragments were used in downstream analysis ^56^.

Differential CUT&RUN enrichments were calculated using pyicos over the previously defined 500bp regions promoter regions. Using bedtools intersect bam all the reads intersecting called peaks were count and used to calculate enrichment fold change with pyicos ^57^ pyicoenrich –counts –pseudocount parameters. Zscore associated p-value and Benjamini Hochberg corrected p-value were computed in R 2*(pnorm(-abs(zscore)) and p.adjust method=BH.

### ABC model

TheABCmodel ^16^wascomputedfollowing https://github.com/broadinstitute/ABC-Enhancer-Gene-Prediction. Processed bam files, as explained above, from bulk ATAC-seq and H3K27ac CUT&Run from Ctr and IFN*γ* treated OPCs were provided as input for the model. Default parameters were used for generating the candidate enhancer list and for quantifying enhancer activity the top 150 000 peaks were considered. Contact frequency was estimated using the power law provided by the model and the powerlaw. Score was used as the final ABC score, with 0.02 as the cutoff threshold.

## Supplementary Materials

**Supplementary Table 1**. Genomic coordinates and associated genes presenting differential chromatin accessibility (scATAC-seq) between EAE and Ctr in OPCs, MOL1/2 and MOL5/6 and Gene Ontology terms for each population, enriched in EAE.

**Supplementary Table 2**. Correlation gene activity score (chromatin accessibility over 500 bp promoter region, from scATAC-seq) with gene expression (from scRNA-seq) in OPCs, MOL1/2 and MOL5/6 and Gene Ontology terms for each type for each population.

**Supplementary Table 3**. List of SNPs and outside variants associated with susceptibility in MS overlapping with chromatin accessibility (from scATAC-seq) in mouse OLG and MiGl in EAE.

**Supplementary Table 4**. Transcription factor (TF) motif enrichment in EAE-OLG populations and the TF RNA expression levels (from scRNA-seq).

**Supplementary Table 5**. Differential gene expression (RNA-seq), chromatin accessibility (500 bp promoter regions and annotated enhancers, from ATAC-seq) and associated Gene Ontology terms between Ctr primary OPCs and OPCs upon IFN*γ* treatment. Correlation gene activity score (chromatin accessibility over 500 bp promoter region, from ATAC-seq) with gene expression (from RNA-seq) between Ctr primary OPCs and OPCs upon IFN*γ* treatment and associated Gene Ontology terms.

**Supplementary Table 6**. Differential gene expression (RNA-seq) between Ctr primary OPCs and OPCs upon Dexamethasone treatment, and associated Gene Ontology terms.

**Supplementary Table 7**. List of genes with differential Cut&Run peaks for H3K27ac, CTCF, H3K27me3 and H3K4me3, between Ctr primary OPCs and OPCs upon IFN*γ* treatment.

**Supplementary Table 8**. List of bivalent genes (marked by both H3K27me3 and H3K4me3) in Ctr primary OPCs.

**Supplementary Table 9**. Differential gene expression (RNA-seq) between EZH2i and Ctr in IFN*γ*- and Ctr-OPCs and associated Gene Ontology terms.

---

**Fig. S1.**
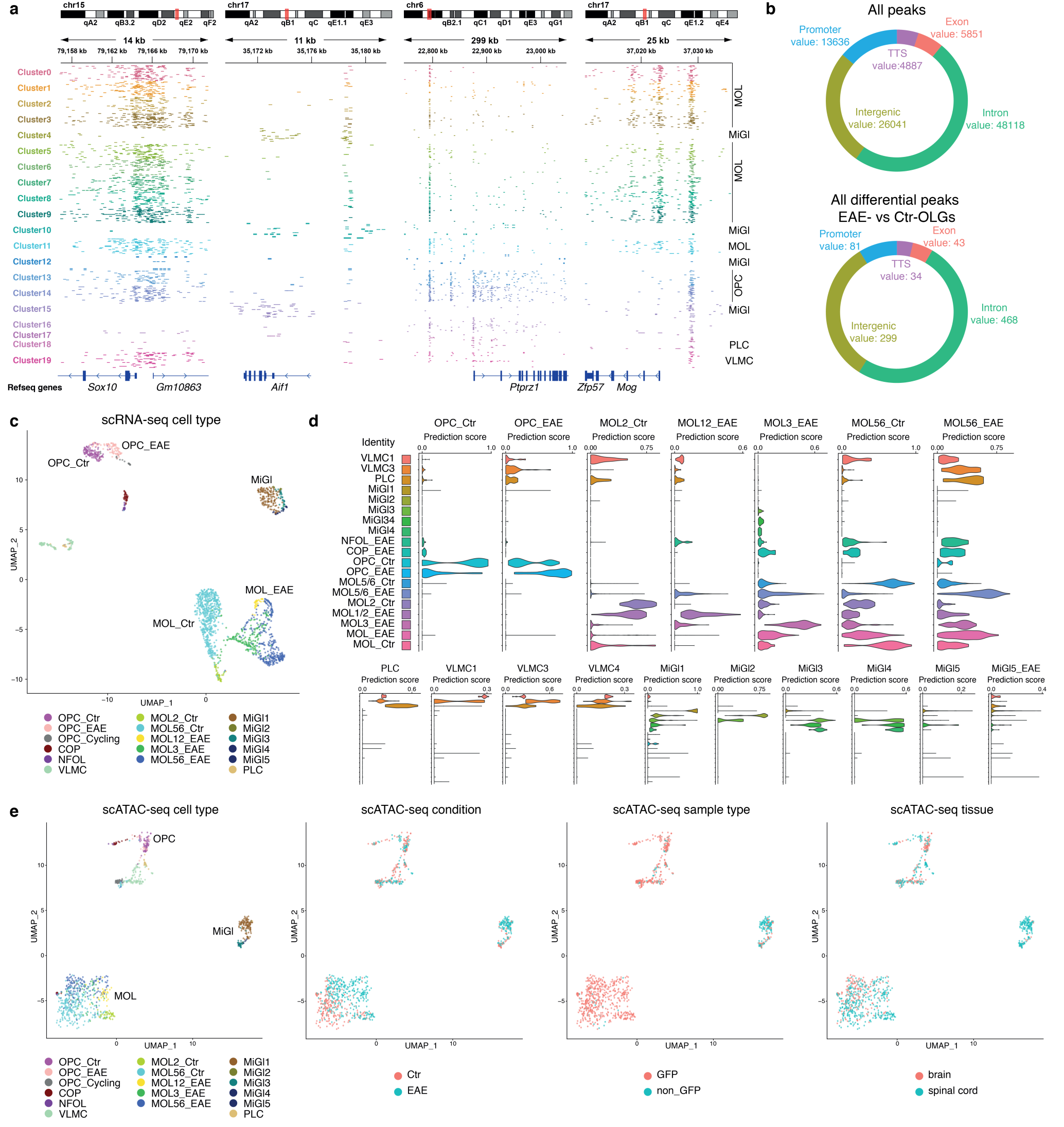
**a**, IGV tracks of chromatin accessibility in 50 randomly selected individual cells for each of 20 cell clusters identified with unsupervised clustering. Colors represent the cluster as in Fig. 1b. Marker genes are shown for OLG (*Sox10*), MiGl (*Aif1*), OPCs (*Ptprz1*) and MOLs (*Mog*). Genomics coordinates are shown. **b**, Distribution of accessibility of all peaks in all cells (All peaks) or differential accessibility peaks between EAE- and Ctr-OLG. TTS = Transcription termination site. **c**, UMAP clustering of scRNA-seq data ^1^, showing the major clusters, colors represent the clusters as in Fig. 1d. **d**, Prediction scores for each population to show the performance of the scRNAseq integration with scATAC-seq data to predict cell-types. **e**, Plate-based scATAC-seq (PI-ATAC) on GFP+ and GFP-cells (Sox10:Cre-RCE:LoxP(EGFP)) collected from brains and spinal cords from EAE and Ctr mice. Clustering performed with UMAP based on differential accessibility. Shown are cell-types (label transfer from scRNA-seq ^1^), condition (EAE vs. Ctr), sample type (GFP vs. non-GFP) and tissue (brain vs. spinal cord).

**Fig. S2.**
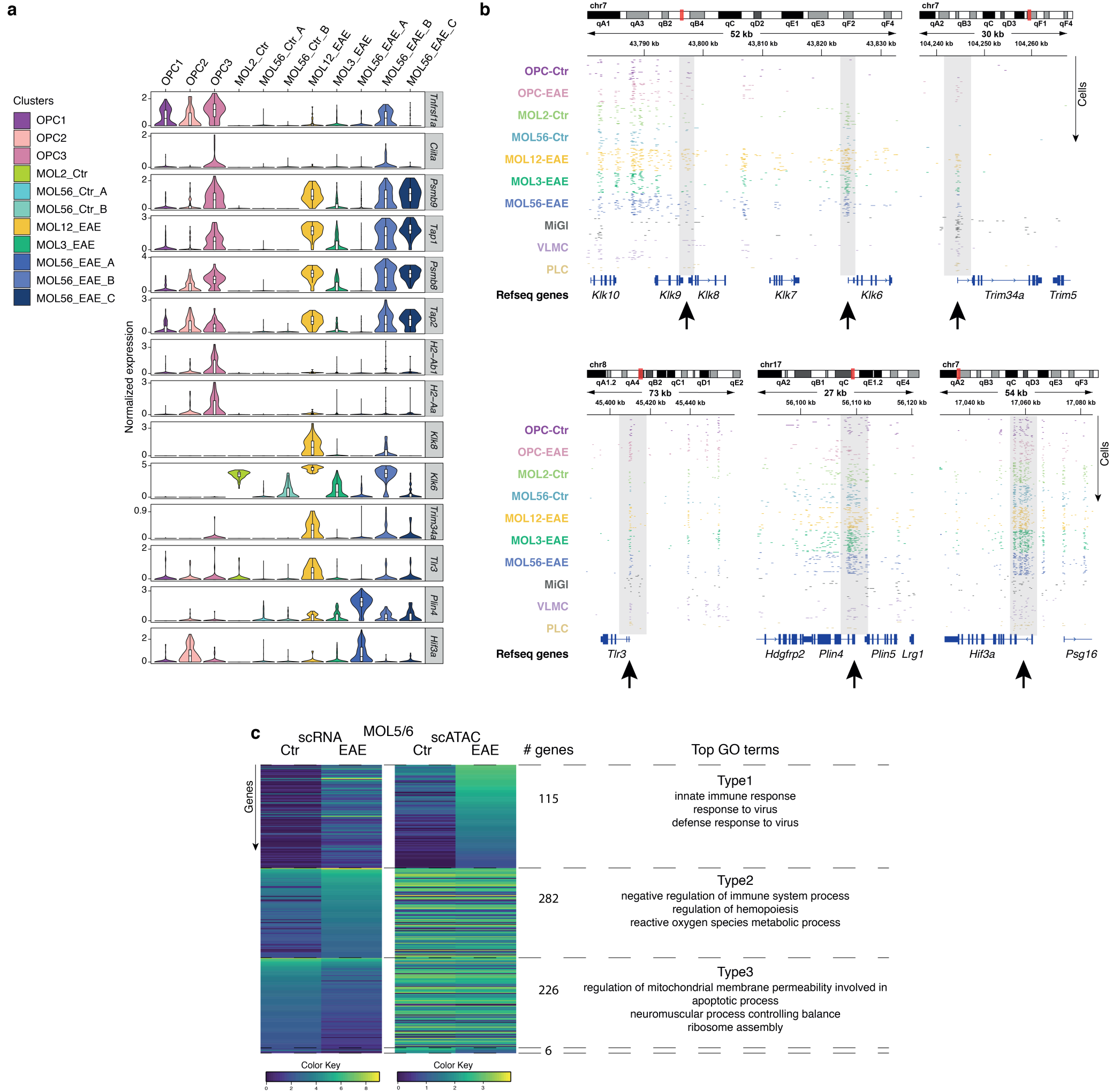
**a**, Violin plots showing normalized RNA expression values per OLG cluster for selected genes (data from ^1^). **b**, IGV tracks of the chromatin accessibility in 50 randomly selected individual cells for each selected cluster, with MiGl clusters grouped together. Highlighted with grey boxes and arrows are regions with differential accessibility in specific clusters or promoter priming, black arrows indicate promoter regions. Genomic coordinates are shown. **c**, Genes in MOL5/6 (EAE vs Ctr) are clustered based on gene expression differences between EAE vs. Ctr and correlation with chromatin accessibility activity score (accessibility over 500 bp promoter region). Top GO terms are shown for Type1 (genes with increased expression in EAE and chromatin accessibility), Type2 (genes with increased expression in EAE, but no change in chromatin accessibility) and Type3 (genes with reduced expression in EAE, but no change in chromatin accessibility). Type4 (genes with reduced expression and chromatin accessibility in EAE) had no GO terms because of too few genes in this group.

**Fig. S3.**
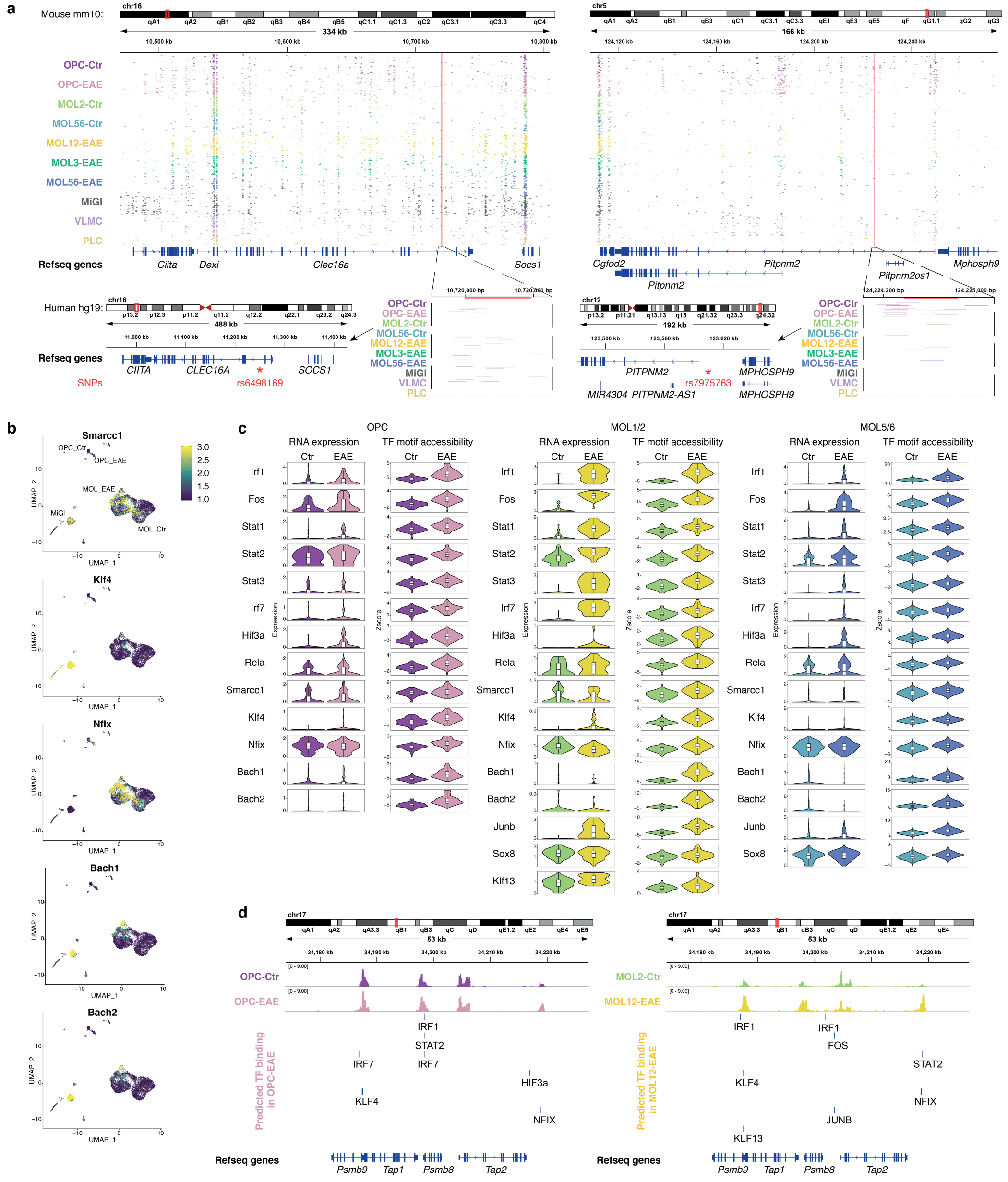
**a**, SNP coordinates for two MS SNPs in the hg19 human genome reference and chromatin accessibility in scATAC-seq in corresponding locations in the mouse mm10 genome reference. IGV tracks of chromatin accessibility in 50 randomly selected individual cells from scATAC-seq are shown. Red boxes show scATAC-Seq peaks from mouse overlapping with SNP location. **b**, TF motif variability projected on top of UMAP clustering. **c**, Violin plots showing RNA expression values of selected TFs and their TF motif accessibility score in OPCs, MOL1/2 and MOL5/6, comparing cells derived from CFA-Ctr and EAE. **d**, IGV tracks of merged single cell chromatin accessibility of different populations and TF predicted binding sites in OPCs and MOL1/2 cells derived from EAE mice in the *Psmb9*-*Tap2* locus.

**Fig. S4.**
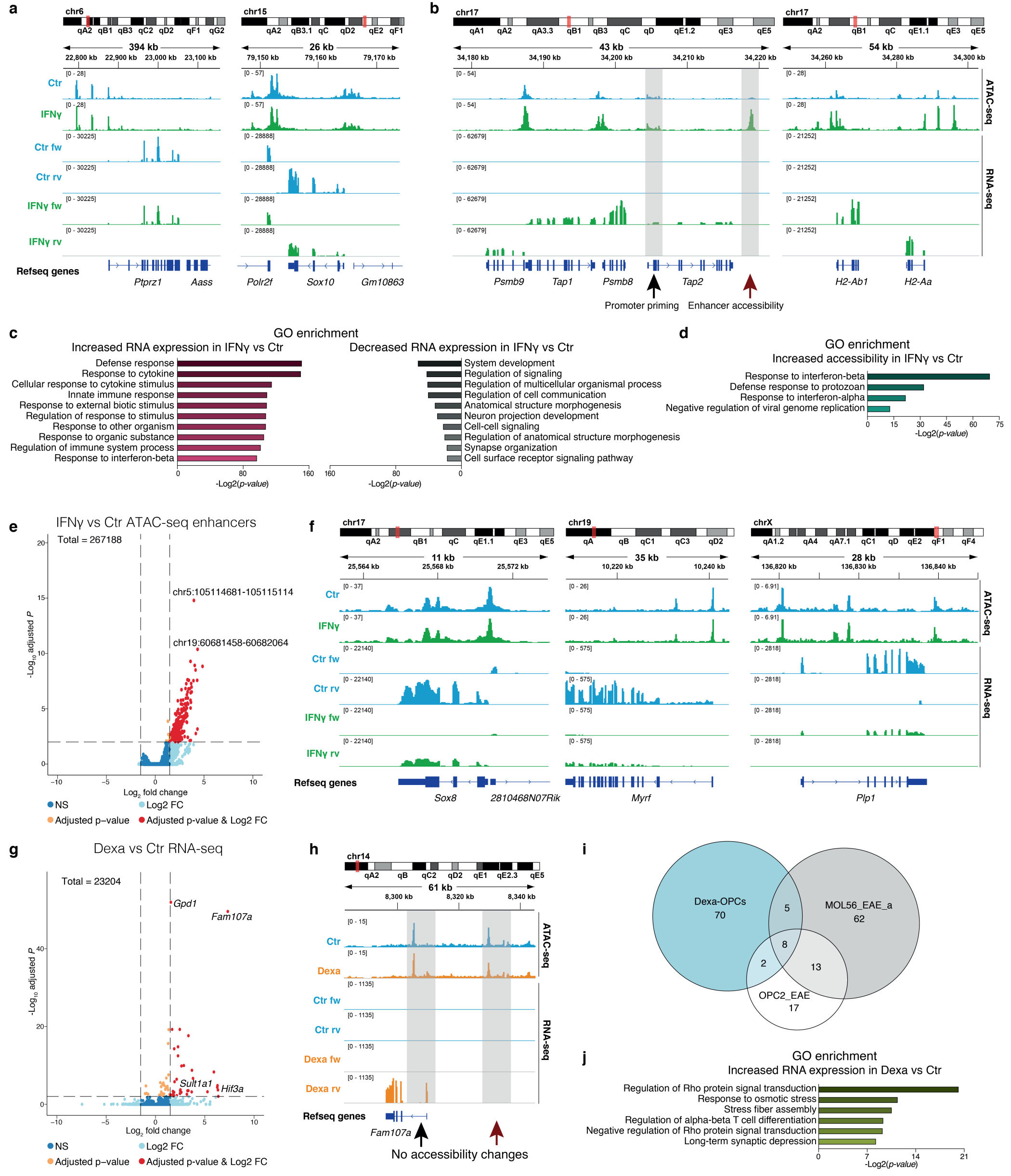
**a, b** IGV tracks of RNA-seq and ATAC-seq from Ctr-OPCs and OPCs treated with 100 ng/ml IFN*γ* for 48 hours, centered at loci for marker genes of OPCs (*Ptprz1*) and OLG (*Sox10*) (**a**) and MHC-I pathway and MHC-II genes (**b**). Highlighted with grey boxes and arrows are regions with differential accessibility in specific conditions or promoter priming, black arrows indicate promoter regions and red arrows indicate putative enhancers. Merged tracks of 3 replicates are shown for ATAC-seq and 4 replicates for RNA-seq. **c**, Top 10 Gene Ontology biological terms for genes upregulated upon IFN*γ* treatment in OPCs and downregulated upon IFN*γ* treatment. **d**, Gene Ontology biological terms for genes with increased chromatin accessibility at 500 bp promoter regions upon IFN*γ* treatment in OPCs. **e**, Volcano plot for chromatin accessibility at annotated enhancers between IFN*γ*-treated and Ctr-OPCs. Loci with statistical significance are shown in orange and genes with statistical significance and log2 fold change above 1.5 are shown in red. **f**, IGV tracks for genes with decreased expression in IFN*γ* treatment. **g**, Volcano plot showing differential gene expression in RNA-seq between Ctr-OPCs and OPCs treated with 1 µm of Dexamethasone (Dexa) for 2 days. Genes with statistical significance are shown in orange and genes with statistical significance and log2 fold change above 1.5 are shown in red. **h**, IGV tracks for a gene with increased expression upon Dexa treatment. Highlighted with grey boxes and arrows are chromatin accessibility regions, black arrows indicate promoter regions and red arrows indicate putative enhancers. **i**, Venn diagram showing the number of genes enriched in Dexa-OPCs vs. Ctr, MOL56-EAE-a vs. MOL56-Ctr-a, OPC2-EAE vs. OPC1 and their overlap between each category. **j**, Gene Ontology biological terms for genes upregulated upon Dexa treatment in OPCs.

**Fig. S5.**
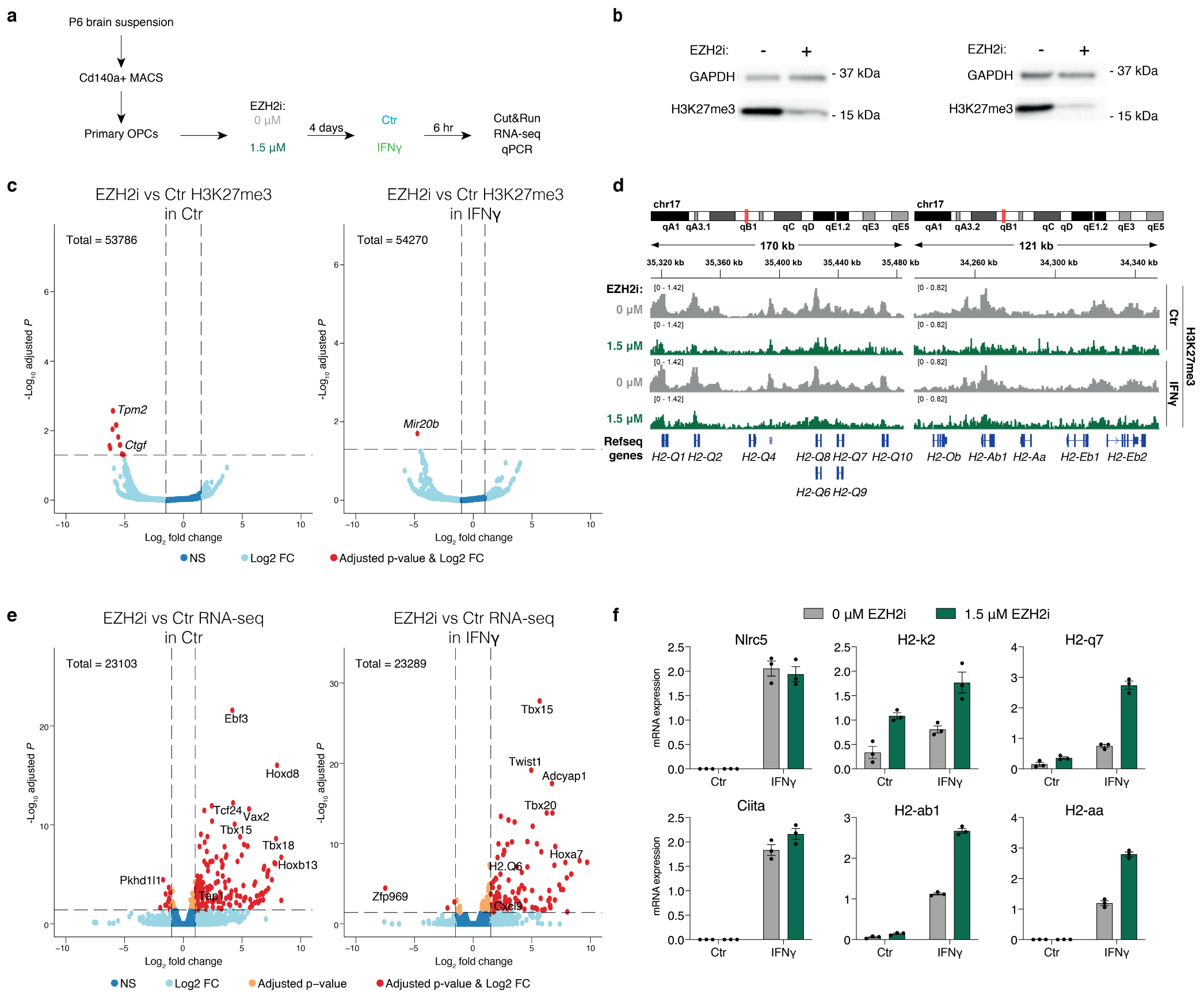
**a**, Schematic overview of EZH2 inhibitor EPZ011989 (EZH2i) experiments. **b**, Western blot with antibodies against H3K27me3 and GAPDH (control) in EZH2i cells vs. Ctr in IFN*γ*-spiked OPCs, two replicates are shown. **c**, Volcano plots for differences in H3K27me3 occupancy between EZH2i vs. Ctr, in Ctr-OPCs (left) and OPCs upon IFN*γ* treatment (right), assessed with Cut&Run. Three replicates are performed. Genes with statistical significance and log2 fold change above 1.5 are shown in red. **d**, IGV tracks for H3K27me3 occupancy upon EZH2i vs. Ctr, in Ctr-OPCs and OPCs upon IFN*γ* treatment at MHC-I and MHC-II loci. Merged tracks of three replicates are shown. **e**, Volcano plots showing differential gene expression between EZH2i vs. Ctr, in Ctr-OPCs (left) and OPCs upon IFN*γ* treatment (right). Genes with statistical significance are shown in orange and genes with statistical significance and log2 fold change above 1 are shown in red. Three replicates are performed. **f**, qRT-PCR analysis of MHC-I and MHC-II pathway genes in OPCs upon treatment with 1,5 µM EZH2 inhibitor EPZ011989 (EZH2i) for 4 days, with subsequent co-treatment with 100 ng/ml IFN*γ* for 6 last hours. Error bars represent SEM, three replicates are shown.

## Notes

https://castelobranco.shinyapps.io/SCATAC10X_2020/

